# UHRF1 upregulation mediates exosome release and tumor progression in osteosarcoma

**DOI:** 10.1101/2020.05.24.113647

**Authors:** Stephanie C. Wu, Ahhyun Kim, Loredana Zocchi, Claire Chen, Jocelyne Lopez, Kelsey Salcido, Jie Wu, Claudia A. Benavente

**Affiliations:** Department of Pharmaceutical Sciences, University of California, Irvine, CA 92697, USA; Department of Developmental and Cell Biology, University of California, Irvine, CA 92697, USA; Department of Biological Chemistry, University of California, Irvine, CA 92697, USA; Chao Family Comprehensive Cancer Center, University of California, Irvine, CA 92697, USA

## Abstract

Loss of function mutations at the retinoblastoma (*RB1*) gene are associated with increased mortality, metastasis and poor therapeutic outcome in several cancers, including osteosarcoma (OS). However, the mechanism(s) through which *RB1* loss worsens clinical outcome remain to be elucidated. Ubiquitin-like with PHD and Ring Finger domains 1 (UHRF1) has been identified as a critical downstream effector of the RB/E2F signaling pathway that is overexpressed in various cancers. Here, we show that UHRF1 upregulation is critical in rendering OS cells more aggressive. Using novel OS genetically engineered mouse models, we determined that knocking-out *Uhrf1* considerably reverses the poorer survival associated with *Rb1* loss. We also found that high *UHRF1* expression correlates with increased clinical presentation of metastatic disease. Based on gain- and loss-of-function assays, we found that UHRF1 promotes cell proliferation, migration, invasion, and metastasis. UHRF1-mediated cell mobility results as a consequence of altered extracellular vesicles and their cargo, including urokinase-type plasminogen activator (uPA). Our work presents a new mechanistic insight into *RB1* loss-associated poor prognosis and a novel oncogenic role of UHRF1 through regulation of exosome secretion that is critical for OS metastasis. This study provides substantial support for targeting UHRF1 or its downstream effectors as novel therapeutic options to improve current treatment for OS.

## INTRODUCTION

Osteosarcoma (OS) is the most common primary cancer of bone and typically occurs in children and young adults. As a highly metastatic cancer, 15-20% of OS patients are diagnosed after the cancer has already metastasized (typically to the lungs), which translates to 5-year survival rates of <40% [1]. In comparison, patients without metastases have survival rates of 65-75%. Unfortunately, pulmonary metastases occur in nearly half of all OS patients [2]. OS survival rates have plateaued since the introduction of adjuvant and neoadjuvant multiagent chemotherapy in the early 1980s [3]. Thus, there is a pressing clinical need to determine the factors responsible for metastasis in OS to facilitate development of new therapeutic strategies.

The most common genetic mutations found in OS are in tumor suppressors *TP53* and *RB1* [4, 5]. Loss-of-function mutations of the *RB1* gene in OS are associated with poor therapeutic outcome as defined by increased mortality, metastasis, and poor response to chemotherapy [6–10]. However, the mechanism(s) through which RB loss leads to poor prognosis remains unclear. A previous study in retinoblastoma identified Ubiquitin-like, containing PHD and RING finger domains 1 (UHRF1) as a key target gene downstream of the RB/E2F signaling pathway that is abnormally upregulated following the loss of RB, and contributes to tumorigenesis [11]. UHRF1 is a multi-domain protein that exerts various functions that include, but are not limited to, reading and writing of epigenetic codes. UHRF1 is most known for its role in maintaining DNA methylation throughout replication by recruiting DNA methyltransferase 1 (DNMT1) to hemi-methylated DNA at the replication fork. In recent years, many reports have emerged showing overexpression of UHRF1 in different types of human cancers that frequently present *RB1* mutations including lung, breast, bladder, colorectal cancer, and retinoblastoma [11–15]. Thus, we hypothesized that loss of RB in OS contributes to tumor progression and increased malignancy through upregulation of its downstream target UHRF1.

In this study, we provide evidence for UHRF1 as a key player in OS tumorigenesis, promoting proliferation, migration, and metastasis. Loss of *Uhrf1* in developmental OS mouse models drastically delayed tumor onset, decreased pulmonary metastasis, and increased the life span of mice carrying *Rb1* mutations. We identified a novel role of UHRF1 in promoting exosome secretion which is associated, at least in part, with the induction of migration and invasion through the secretion of proteolytic enzymes including urokinase-type plasminogen activator (uPA) to remodel the extracellular matrix. Targeting uPA with small-molecule inhibitors resulted in a robust decrease in OS migration and invasion. These findings highlight a critical role of UHRF1 as a driver of the poor prognosis associated with RB loss and presents UHRF1 and its downstream targets as a novel therapeutic targets for OS.

## MATERIALS AND METHODS

### Xenografts, mouse models and cell lines

The five orthotopic xenografts used in this study: SJOS001105 (PDX1), SJOS001112 (PDX2), SJOS001107 (PDX3), SJSO010930 (PDX4), and SJOS001121 (PDX5) were obtained from the Childhood Solid Tumor Network [16]. Athymic nude (NU/J) mice were obtained from The Jackson Laboratories. The *p53^lox^* and *Rb1*^lox^ conditional-knockout (cKO) mice were obtained from the Mouse Models of Human Cancer Consortium at the National Cancer Institute; the *Osx-cre* mice were obtained from The Jackson Laboratory. For *Uhrf1* cKO mice, mice obtained from the European Mouse Mutant Archive were backcrossed to flipasse (Flp) mice for removal of the neo-cassette (tm1c conversion), and then backcrossed to C57BL/6N mice for Flp removal. Mice were monitored weekly for signs of OS. Moribund status was defined as the point when tumors had reached 10% body weight or induced paralysis in the mouse. The University of California Irvine Institutional Animal Care and Use Committee approved all animal procedures.

OS cell lines 143B, SJSA-1, SaOS-2 and U-2 OS, as well as MSCs and HEK293T cells were acquired from ATCC and maintained according to their specifications.

### FDG-PET/microCT scan

Overnight-fasted mice were injected with 0.1-0.5 mCi of ^18^F-FDG in sterile saline (0.05-0.2 ml) intraperitoneally (i.p.) and allow to uptake the ^18^F-FDG for 60 min prior to imaging. Animals were anesthetized and laid in supine position on the scanner holder with continued anesthesia. Scanning data was acquired in full list mode and sorted into a single frame, 3 dimensional sinogram, which was rebinned using a Fourier rebinning algorithm. The images were reconstructed using 2-dimensional filter back projection using a Hanning Filter with a Nyquist cut off at 0.5 and corrected for attenuation using the Co-57 attenuation scan data. Analysis of PET was conducted using PMOD 3.0 and IRW software. The PET data was co-registered to the CT template for drawing regions-of-interest (ROI). The ROI data was converted to standard uptake value (SUV) of ^18^F-FDG.

### Western blotting

Western blots were performed as described [17] using the following antibodies and dilutions: 1:1000 anti-Uhrf1 (sc-373750, Santa Cruz Biotechnology), 1:1000 anti-Rb (9313T, Cell Signaling), anti-actin (A1978, Sigma), anti-E2F1 (3742S, Cell Signaling), anti-E2F2 (sc-9967, Santa Cruz Biotechnology), anti-E2F3 (MA5-11319, Nalgene Nunc). Secondary antibodies were diluted 1:1000 (PI-1000 or PI-2000, Vector Laboratories). Bands were visualized using chemiluminescence (SuperSignal™ West Pico Chemiluminescent Substrate, Thermo Scientific). Band intensities were analyzed using ImageJ software.

### *In situ* hybridization

OS tissue microarrays were purchased from US Biomax, Inc (OS804c). 6 stage IA (T1N0M0), 16 stage IB (T2N0M0), 1 stage IIA (T1N0M0), 16 stage IIB (T2N0M0) and 1 stage IVB (T2N1M0) tumors are represented in the array. Stage I tumors are low grade and stage II and IV are high grade. The extent of the primary tumor (T) is classified as intercompartmental (T1) or extracompartmental (T2). Tumors that have metastasized to nearby lymph nodes are denoted by N1, but no distant organ metastases (M0) are represented in the array. Formalin-fixed, paraffin-embedded (FFPE) slides were incubated at 60°C for 1hr for deparaffinization, followed by two washes of xylene and two washes of 100% ethanol for dehydration, 5 min each at RT, before air-drying. RNAscope^®^ technology was utilized for *in situ* hybridization. Assay was performed following manufacturer’s protocol. Human and mouse UHRF1-specific probes were customized by Bio-Techne. RNAscope^®^ Positive Control Probe- Hs-PPIB/Mm-PP1B and RNAscope^®^ Negative Control Probe- DapB were used. OpalTM 570 fluorophore was used at a 1:1000 dilution. Slides were mounted with ProLong Gold Antifade Mountant.

### Chromatin immunoprecipitation (ChIP assay)

ChIP assays were performed as previously described [18]. ChIP DNA was analyzed by qPCR with SYBR Green (Bio-Rad) in ABI-7500 (Applied Biosystems) using the following primers. For motif 1 (M1), Forward: 5’-CCACATTCCCTCGCAGTATTTA-3’; Reverse 5’-CCCTGAACTCTTAAGTCCAAGTC-3’; for motifs 2 and 3 (M2, M3), Forward: 5’-CACCCTCTTTCTCGCTTCC-3’; Reverse 5’-TGCCAGCTGCTCTGATTT-3’. The anti-E2F1 (3742; Cell Signaling) and rabbit IgG (sc-2027, Santa Cruz Biotechnologies) antibodies were used.

### Real-time RT-PCR (qPCR)

We performed qPCR as described [17]. Primers were designed using IDT Real-Time PCR tool (Integrated DNA Technologies). Reaction was carried out using 7500 Real-Time PCR system (Applied Biosciences). Data were normalized to those obtained with endogenous control 18S mRNA and analyzed using ΔΔCt method. Primer sequence for PCR amplification are as follows: *UHRF1* (5’-GCTGTTGATGTTTCTGGTGTC-3’; 5’-TGCTGTCAGGAAGATGCTTG-3’), *PLAU* (5’-GAGCAGAGACACTAACGACTTC-3’; 5’-CTCACACTTACACTCACAGCC-3’), *SEMA3E* (5’-CTGGCTCGAGACCCTTACTG-3’; 5’- CAAAGCATCCCCAACAAACT-3’), *HAS2* (5’-TCCATGTTTTGACGTTTGCAG- 3’; 5’-AGCAGTGATATGTCTCCTTTGG-3’), *LAMC2* (5’-CACCATACTCCTTGCTTCCTG-3’; 5’- GTGCAGTTTGTCTTTCCATCC-3’), *18S* (5’-GTAACCCGTTGAACCCCATT-3’; 5’-CCATCCAATCGGTAGTAGCG-3’).

### Lentivirus production and transduction

Lentiviral particles were produced by co-transfecting the envelope plasmid pCMV-VSV-G (Addgene), packaging plasmid pCMV-dR8.2 dvpr (Addgene), lentiCRISPRv2 (Addgene) or plentiCRISPRv2 (GenScript) with UHRF1 gRNA (gRNA sequence: TCAATGAGTACGTCGATGCT), inducible CRISPR/Cas9 plasmid TLCV2 (Addgene plasmid #87360) with UHRF1 gRNA (gRNA1 sequence: CGCCGACACCATGTGGATCC and gRNA2 sequence: ACACCATGTGGATCCAGGTT) into HEK293T cells using calcium phosphate transfection method. Supernatants containing lentiviral particles were harvested at 24 h and 48 h post-transfection. Cell debris were cleared by centrifugation at 1600 × g for 10 min at 4°C. Supernatants were then filtered through 0.45 μm PES filter (25-223, Genesee Scientific), and concentrated by ultracentrifugation at 23000 RPM for 2 h at 4°C. Lentiviral particles were resuspended in ice-cold PBS and stored at −80°C. Transduction of target cells were achieved by exposing cells to viral particles in serum-free condition for 6 hr. Puromycin selection was carried out at a concentration of 2 μg/ml.

### Clonogenic assay

6-well plates were coated with 1ml of 0.75% agarose prior to seeding. Single cell suspension was mixed with agarose to achieve a final agarose concentration of 0.36%, the mixture was then layered on top of the 0.75% agarose coating (Seeding density 200 cells/cm^2^). Cells were incubated at 37°C with 5% CO_2_ until cells in control wells have formed sufficiently large colonies (>50 cells). Colonies were fixed with 10% methanol for 15 min and stained with 5% Giemsa for 30 min for visualization.

### Immunocytochemistry

Cell were seeded onto glass coverslips in a 24-well plate at a density of 25000 cells/well and allowed to attach overnight. For EdU staining the Click-iT EdU cell proliferation kit (Thermo Fisher Scientific) was used following manufacturer’s instructions with 1 h pulse using a final 10 μM EdU solution in media. For UHRF1 immunostaining, media was aspirated, and cells were fixed using 4% (w/v) paraformaldehyde overnight at 4°C. Slides washed with PBS twice and then incubated with primary antibody overnight at 4°C. Mouse anti-UHRF1 antibody (PA5-29884, Thermo Fisher Scientific) was used at 1:500 dilution. Slides washed 3 times in PBS then incubated with anti-rabbit secondary antibody (BA-1000, Vector Laboratories) at 1:500 dilution with corresponding blocking buffer for 30 min at RT in the dark. Slides were washed 3 times with PBS and incubated in 300 μl of Vectastain ABC kit (Vector Laboratories) for 30 min at RT. Slides were washed 3 times in PBS and incubated with tyramide Cy3 1:150 in amplification buffer (Perkin Elmer) for 10 min. After washing with PBS for 3 times, DAPI was added (1:1000 dilution in PBS) for 5 min to stain the nuclei. Slides were washed twice with PBS and mounted using gelvatol containing DABCO. Imaging was completed using Life Technologies EVOS microscope. Cell scoring was performed by a blinded investigator.

### Subcutaneous injection

Human OS cell lines were dissociated in 0.25% trypsin-EDTA, washed with PBS prior to counting. 2·10^6^ cells were resuspended in 100 μl of PBS and injected subcutaneously into the flank region of mice. Tumors were collected 3 weeks post injection. For cells that require doxycycline induction for the expression CRISPR/Cas9, 2 mg/ml doxycycline hyclate supplemented water was administered *ad libitum*. 10 mg/ml of sucrose was added to doxycycline supplemented water to increase consumption.

### RNA Sequencing

Total RNA was isolated using the RNA Micro Kit (Qiagen). Subsequently, 500 ng of total RNA was used to create the RNA-seq library following the manufacturer’s protocol from purification, mRNA fragmentation through the adenylation of end-repaired cDNA fragments and cleanup (TruSeq Stranded mRNA, Illumina). The collected sample was cleaned with AMPure XP beads (Beckman Coulter) and eluted in 20 μl of 10 mM Tris buffer, pH 8, 0.1% Tween 20. A paired-end 100-bp sequencing run was performed on HiSeq 4000 yielding 348M PE reads with a final library concentration of 2 nM as determined by qPCR (KAPA Biosystem).

Sequencing reads were aligned to the mouse or human reference genome (GRCm38 or GRCh38, ENSEMBL v.92) using STAR (v2.5.2a) [19]. Each read pair was allowed a maximum number of mismatches of 10. Each read was allowed a maximum number of multiple alignments of 3. Gene counts for each sample were produced using HTSeq (v0.6.1p1) [20]. Read count normalization and differential expression analysis were performed using DESeq2 (v1.22.2) in R (v3.5.2) [21]. Genes with low reads (sum across samples < 10) were removed. Differentially expressed genes (DEG) were calculated by comparing UHRF1 knockouts to matching controls, genes with base mean ≥ 10, Log2FoldChange ≥ 1 or ≤ −1, and adjusted p value (Benjamini-Hockberg) ≤ 0.05 were called differentially expressed genes. 3D PCA plot was generated using the R package plot3D (v1.1.1).

All sequencing data discussed in this publication have been deposited in NCBI’s Gene Expression Omnibus [22] and are accessible through GEO Series accession number GSE144418 (https://www.ncbi.nlm.nih.gov/geo/query/acc.cgi?acc=GSE144418).

### Scratch-wound assay

Cells were seeded the day before the scratch at a density of 1.1·10^5^/cm^2^ and grown to 100% confluency in 6-well cell culture plates. 2 h before the scratch, cells were treated with 5 μg/ml of mitomycin C (S8146, Sigma). At time 0, a 1000 μl tip was used to create a wound across the cell monolayer. The cells were incubated for 8 h at 37°C with 5% CO_2_. Images of the wound at the exact same field were taken at 0 and 8 h and analyzed using the ZEISS ZEN microscope software. A total of 20 measurements in pixels were made per image. The pixels migrated was calculated as the difference between the average width of the scratches before and after the incubation period. p-values were calculated using two-tailed t-test. Amiloride was used at 150 μM, BC 11 hydrobromide at 16.5 μM, GW4869 at 10 μM, which were supplemented in growth medium and given to the cells at the 0 h time point. Chlorpromazine (CPZ) was used at 20 μM for 30 min prior to scratch. For experiments using conditioned media, serum-free media was used and exposed to cells for 24 h.

### Transwell invasion assay

8 μm pore PET membranes (353097, Corning) were coated with Matrigel (25 μg/insert). 2·10^4^ cells were seeded onto the membrane in 100 μl of appropriate cell culture medium and placed into the incubator for 2 h at 37°C with 5% CO_2_ to allow cell attachment. After cells have attached, 100 μl of media were added to the inner chamber, amiloride (150 μM final concentration) or BC11 hydrobromide (16.4 μM final concentration) or doxycycline (1 μg/ml final concentration) were supplemented at this time. 600 μl of appropriate cell culture medium supplemented with 100 ng/ml fibroblast growth factor (FGF) was added to the outer chamber. Cells were incubated for 16 h at 37°C with 5% CO_2_. Cell culture medium was aspirated, and cotton swabs were used to scrape off non-migratory cells on the top side of the insert. The remaining cells at the bottom side of the insert were then fixed and stained with a solution consists of 6% glutaraldehyde and 0.5% crystal violet for 2 hr. Stains were washed out with tap water and the inserts left to dry at room temperature. Images of 5 different fields within each insert were taken for analysis by cell count.

### Tail vein injection

Human OS cells lines were dissociated in 0.25% trypsin-EDTA, washed with PBS prior to counting. For the experimental metastasis model, 2×10^6^ cells were resuspended in 200 μl of PBS and injected intravenously through the tail vein. Mice lungs were collected 3 weeks after injection, fixed in 4% formaldehyde for histological analysis.

## RESULTS

### UHRF1 overexpression in mouse OS is a critical driver of the poor survival in *Rb1*-null OS

The OS developmental mouse model using Osterix-driven Cre-recombinase (Osx-cre) to facilitate preosteoblast-specific loss of *Tp53* results in OS tumor formation with complete penetrance and gene expression profiles, histology, and metastatic potential comparable to that of human OS [23, 24]. Loss of *Rb1* in addition to *Tp53* potentiates OS in these mice, strongly mimicking the behavior of the disease in human. These OS mice develop tumors with calcified interior and highly proliferative exterior, which can be captured by microCT and FDG-PET scans, respectively (Fig. 1A). Our previous work identified *UHRF1* as a target overexpressed upon *RB1* inactivation that plays a significant role in the epigenetic-driven tumor progression in retinoblastoma [11]. Analysis of UHRF1 expression in OS tumors arising in *Tp53* single conditional-knockout (cKO; *Osx-cre Tp53*^lox/lox^) and *Tp53*/*Rb1* double conditional-knockout (DKO; *Osx-cre Tp53*^lox/lox^ *Rb1*^lox/lox^) OS tumors revealed UHRF1 is highly expressed both at the mRNA (Fig. 1B) and protein level compared to mouse mesenchymal stem cells (mMSC; Fig. 1C).

**Figure 1.**
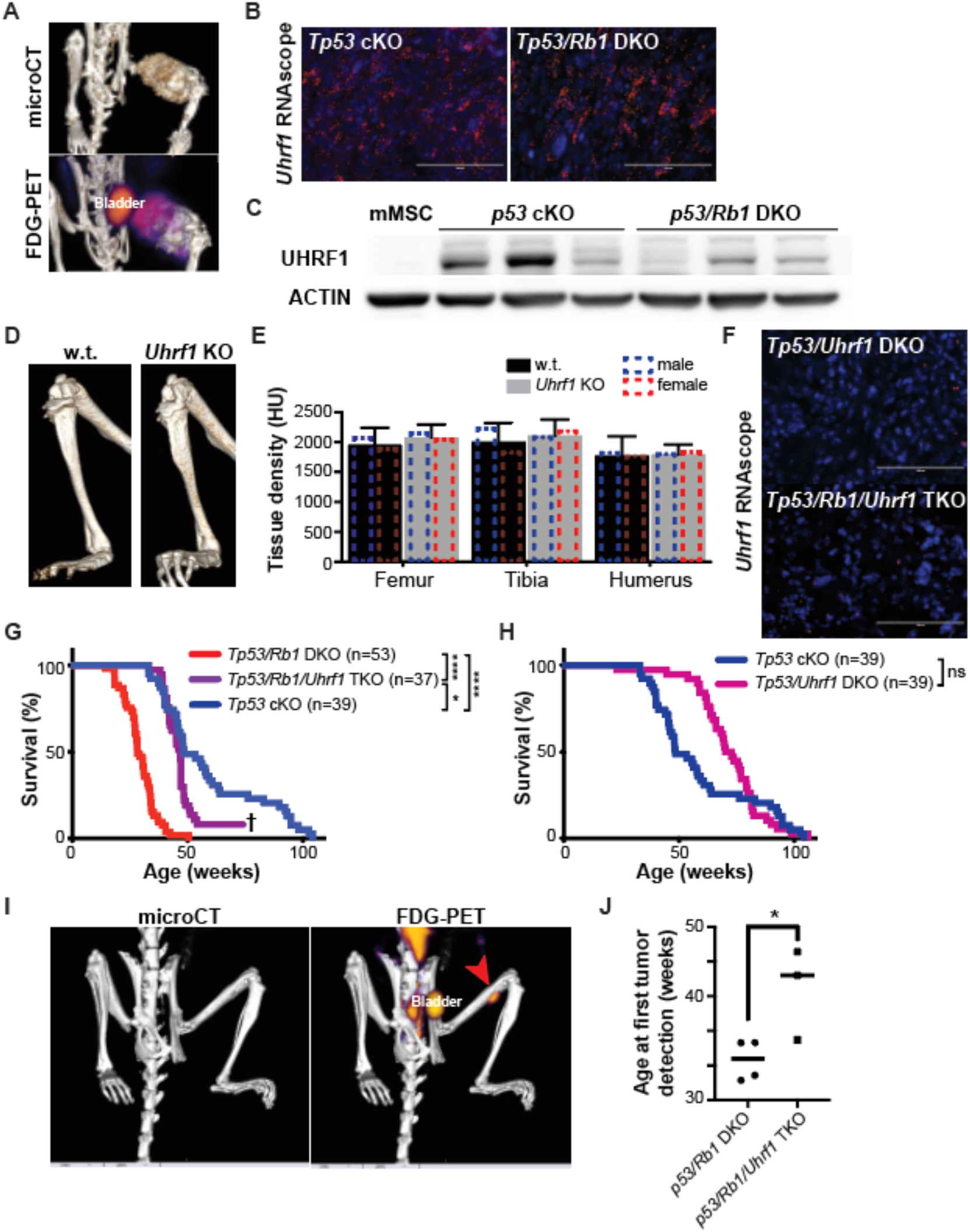
UHRF1 is a critical driver of the increased malignancy observed in *Rb1* null OS. (A) Representative micro-CT (top) and micro-PET (bottom) scans of OS that arises in a mouse femur. (B) Representative fluorescent images of *in situ* hybridizations using RNAscope on tumors from genetically engineered OS mice using a probe against *Uhrf1* (red). Nuclei were counterstained with DAPI (blue). *Uhrf1* expression is detected in *Tp53* cKO and *Tp53/Rb1* DKO mice. (C) Western blot detection of UHRF1 in tumors from genetically engineered *p53* cKO and *p53/Rb1* DKO OS mice, β-actin was used as loading control. (D) Micro-CT scan of hind limb from wildtype (w.t.) and preosteoblastic *Uhrf1* knockout (*Uhrf1* KO) mice (E) Bone density analysis of prime OS locations in Hounsfield units (HU). Each data point is mean ± s.d. of n=4 samples. (F) Representative fluorescent images of *in situ* hybridizations using RNAscope on tumors from genetically engineered OS mice using a probe against *Uhrf1* (red). Nuclei were counterstained with DAPI (blue). *Uhrf1* expression is not detected in *Tp53/Uhrf1* DKO and *Tp53/Rb1/Uhrf1* TKO mice. (G-H) Kaplan-Meier curves showing the survival of OS-prone mouse models. (G) Mice bearing *Rb1* mutations *Tp53/Rb1* DKO: *Osx-Cre; p53^lox/lox^; Rb1^lox/lox^* (red; n=53) have significantly shorter lifespan compared to *Tp53* cKO: *Osx-Cre; p53^lox/lox^; Rb1^lox/lox^* (blue; n=39). This survival time was significantly increased in *Tp53*/*Rb1*/*Uhrf1* TKO: *Osx-Cre; p53^lox/lox^; Rb1^lox/lox^*;*Uhrf1^lox/lox^* (purple; n=37) mice. † Three surviving mice were removed from study to confirm tumor absence. (H) *Tp53*/*Uhrf1* DKO: *Osx-Cre; p53^lox/lox^; Uhrf1^lox/lox^* (magenta; n=39) showed an overall survival comparable to *Tp53* cKO (blue; n=39). Mantel-Cox test were used for curve comparisons. ns=not significant, * p < 0.05, **** p < 0.0001. (I) Representative images from micro-CT (left) and micro-PET (right) scans used for determining the age at which tumors can be first detected (red arrow). (J) Age of mice (in weeks) at earliest tumor detection via micro-CT and PET scans. Solid line represents the average for *Tp53/Rb1* DKO (n=4) and *Tp53*/*Rb1*/*Uhrf1* TKO (n=3). *p < 0.05 by unpaired two-tailed *t* test.

Before examining the role of UHRF1 in OS development *in vivo*, we first assessed the effect of *Uhrf1* loss in normal bone development. Since *Uhrf1* KO mice result in mid-gestational lethality [25], we generated *Uhrf1* cKO mice (*Osx-cre Uhrf1*^lox/lox^). Bone density analyses of the femur, tibia, and humerus exhibited no difference in Hounsfield units (n=4) upon preosteoblastic loss of *Uhrf1* in either male or female mice (Fig. 1D-E). Thus, *Uhrf1* loss does not appear to affect normal bone development, independent of sex (Fig. 1E). Then, we crossed our *Uhrf1* cKO with the developmental OS mouse models *Tp53* cKO and *Tp53*/*Rb1* DKO, resulting in *Tp53*/*Uhrf1* DKO (*Osx-cre Tp53*^lox/lox^ *Uhrf1*^lox/lox^) and *Tp53*/*Rb1*/*Uhrf1* triple knockout (TKO; *Osx-cre Tp53*^lox/lox^ *Rb1*^lox/lox^ *Uhrf1*^lox/lox^). Overall survival and tumor formation of *Tp53*/*Uhrf1* DKO and *Tp53*/*Rb1*/Uhrf1 TKO mice were tracked and compared to their corresponding littermate controls (*Tp53* cKO and *Tp53/Rb1* DKO, respectively) to assess the role of UHRF1 in osteosarcomagenesis and metastasis. All genotypes analyzed led to the development of OS in mice, but with distinct survival rates and disease presentation. *In situ* hybridization performed on the OS mouse tumors confirmed a significant reduction of *Uhrf1* in *Tp53*/*Uhrf1* DKO and *Tp53*/*Rb1*/*Uhrf1* TKO (Fig. 1F). In line with previous reports, the loss of *Rb1* in *Tp53*/*Rb1* DKO mice resulted in a significantly shorter median survival (28.1 weeks; n=53) compared to *Tp53* cKO mice (48.4 weeks; n=39, p<0.0001), recapitulating the poor prognosis associated with *RB1* loss in humans (Fig. 1G). Strikingly, genetic ablation of *Uhrf1* in *Tp53/Rb1/Uhrf1* TKO mice resulted in a significant increase in survival compared to *Tp53/Rb1* DKO mice, with a median survival of 46.9 weeks (n=37; p<0.0001). The median survival of *Tp53/Rb1/Uhrf1* TKO (46.9 weeks) was comparable to that of *Tp53* cKO (48.4 weeks, p=0.0577, Gehan-Breslow-Wilcoxon test), suggesting that elevated UHRF1 is critical for the poor prognosis associated with *Rb1* loss (Fig. 1G). However, the overall survival of *Tp53/Rb1/Uhrf1* TKO and *Tp53* cKO mice is distinct from each other (p=0.0126, Mantel-Cox test), suggesting that early UHRF1 overexpression is not the only factor contributing to *Rb1* loss-associated outcomes. Since three *Tp53/Rb1/Uhrf1* TKO mice had not acquired tumors well beyond the average of the rest of their study group (> 69 weeks; Fig. 1G†), we performed FDG-PET/microCT scans on these mice, followed by autopsy. FDG-PET/microCT scans revealed no detectable tumors (Supp. Fig. 1), which was further confirmed through autopsy, suggesting that *Uhrf1* loss likely prevents early stages of tumor initiation or tumor progression in this OS model. Most remarkably, genetic ablation of *Uhrf1* also resulted in a significant reduction in pulmonary metastatic potential, with only 10% of mice presenting lung metastases in *Tp53/Rb1/Uhrf1* TKO compared to 40% in *Tp53/Rb1* DKO mice. In addition to the reduced metastatic incidence rate, a lighter burden of metastatic nodules was also observed in *Tp53/Rb1/Uhrf1* TKO mice.

Since increased *Uhrf1* mRNA and protein levels were also detected in *Tp53* cKO mouse tumors (Fig. 1B-C), we examined the effect of *Uhrf1* loss in *Tp53/Uhrf1* DKO compared to *Tp53* cKO mice. We observed a significant increase in the median survival of *Tp53/Uhrf1* DKO mice (71.4 weeks; n=39), compared to *Tp53* cKO (n=39; p=0.0007, Gehan-Breslow-Wilcoxon test), but not in overall survival (p=0.1189, Mantel-Cox test; Fig. 1H). Interestingly, we found that all *Tp53* cKO mice analyzed with early morbidity (< 50 weeks) presented tumors with increased *Uhrf1* mRNA levels, suggesting spontaneous RB pathway inactivation, when compared to tumors from mice that reached moribund status later in life (> 50 weeks; Supp. Fig. 1B). This increased *Uhrf1* mRNA expression correlated with increased *Cdk4* expression in these tumors (Supp. Fig. 1C). No correlation was found with the expression of other RB regulators, including *p16* and *Cdk6*, even though the only tumor with undetectable *p16* expression was observed a young tumor (Supp. Fig. 1D-E). No significant changes on *Rb1* expression were observed between these groups (Supp. Fig. 1F). Pathology analysis of histological sections for tumors from the four OS mouse models analyzed linked *Uhrf1* loss with lower mitotic index and decreased anaplasia (Supp. Fig. 2 and Supp. Table 1).

The observation that UHRF1 loss significantly increased overall survival in *Tp53/Rb1/Uhrf1* TKO but not *Tp53/Uhrf1* DKO, along with the observation of three *Tp53/Rb1/Uhrf1* TKO mice with undetectable tumors, suggested that early RB-pathway inactivation, and the associated UHRF1 overexpression, may contribute to early stages of tumor promotion. To determine whether *Uhrf1* contributes to tumor promotion or tumor progression, we performed periodic FDG-PET/microCT scans on *Tp53/Rb1* DKO and *Tp53/Rb1/Uhrf1* TKO mice to determine the age at which tumors are detectable (Fig. 1I). In *Tp53/Rb1* DKO mice, tumors were detected in average at 30.8 ± 2.9 weeks (n=4) while in *Tp53/Rb1/Uhrf1* TKO mice, tumors were detected significantly later at 41.1 ± 6.6 weeks (n=4; p=0.0362; Fig. 1J). All test subjects reached humane end-of-study within 3 weeks of first tumor detection. These data suggest that *Uhrf1* plays a pivotal role in the early developmental process of OS tumor promotion following loss of *Rb1*.

### UHRF1 overexpression correlates with increased malignancy and metastasis in human OS

Following the observation of UHRF1 contributing to a more aggressive phenotype in developmental mouse models of OS, we next examined the role of UHRF1 in human OS. Upon examination of The Cancer Genome Atlas (TCGA) database, we found that *UHRF1* is significantly overexpressed in malignancies that frequently inactivate the RB-pathway including sarcomas, breast invasive carcinomas, and lung adenocarcinomas; while tumors where RB loss is infrequent in primary disease, such as prostate adenocarcinomas [26], show no significant difference in *UHRF1* compared to control normal tissue (Fig. 2A). Furthermore, in sarcomas, high *UHRF1* expression is associated with poorer survival (Fig. 2B). Since OS is not represented in the TCGA sarcoma tumor cohort, we performed *in situ* hybridization of a human OS tissue array to identify the presence of *UHRF1* mRNA in the 25 primary OS tumors examined (Fig. 2C). We observed that tumors expressing high *UHRF1* mRNA levels were classified as stage II and none as stage I OS (Fig. 2D), suggesting that UHRF1 expression increases in more aggressive tumors. However, the limited number of stage III and stage IV samples available restricted our analysis of metastatic cases. Given the interesting possibility of a role of UHRF1 in OS metastasis shown by our mouse models, we analyzed *UHRF1* expression from a published RNA-Seq dataset of pretreatment biopsies from 53 high-grade OS patients who did or did not develop metastasis within 5 years of initial diagnosis, compared to normal mesenchymal stem cells (MSC), and osteoblasts [27]. *UHRF1* expression in MSCs (log2 median = 9.055) was similar to osteoblasts (log2 median = 8.510; p=0.1495) and tumors from patients that did not develop metastasis within 5 years from the time of biopsy (log 2 median = 9.055; p=0.1123). However, tumors from patients who developed metastasis within 5 years from the time of biopsy expressed significantly higher levels of *UHRF1* (log2 mean = 9.880; p = 0.0015; Fig. 2E).

**Figure 2.**
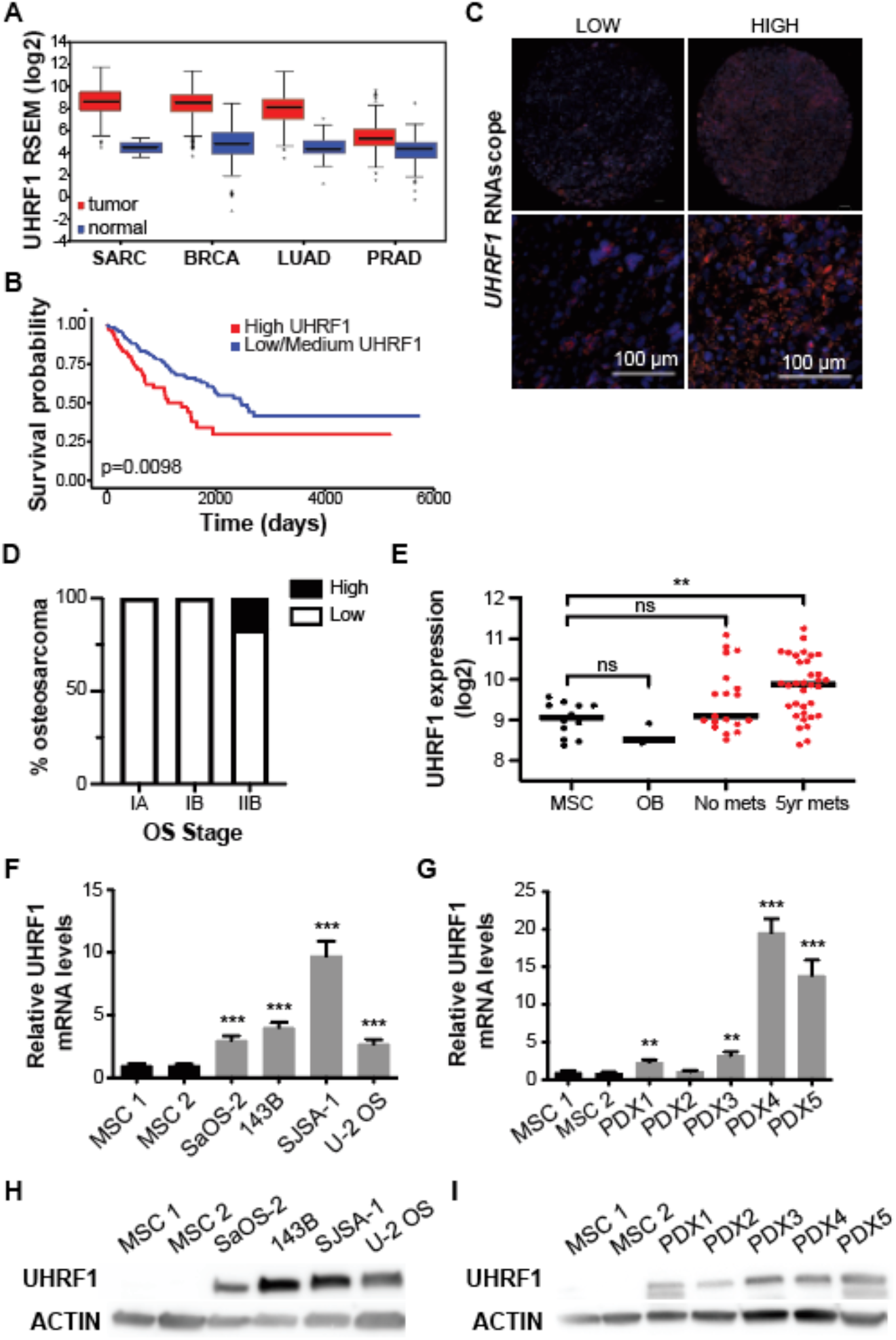
UHRF1 is upregulated and overexpressed in human OS. (A-B) The Cancer Genome Atlas data of (A) RNA-Seq by expectation-maximization (RSEM) values from 263 human sarcoma (SARC) tumor samples compared to 2 normal tissue (17.9-fold), 1100 breast invasive carcinoma (BRCA) compared to 112 normal (13.5-fold), 517 lung adenocarcinoma (LUAD) compared to 59 normal (13.7-fold) show increased *UHRF1* expression in tumors with frequent RB pathway alterations but not in 498 prostate adenocarcinoma (PRAD) tumors compared 52 normal (1.84-fold) were RB pathway is normally unaltered. Tumor samples (red) compared to normal samples (blue). (B) Effect of *UHRF1* expression level on SARC patient survival comparing high UHRF1 expression (red; n=65) to low/medium UHRF1 expression (blue; n=194). (C) Representative fluorescent images in 2X (top) and 20X (bottom) magnification of *in situ* hybridizations using RNAscope scored as low or high based on signal intensity for *UHRF1* mRNA. Scale bar = 100 μm. (D) Quantification of the percentage of low and high signal score acquired from RNAscope in stage IA, IB, and IIB OS. (E) RNA-seq analysis comparing *UHRF1* expression between mesenchymal stem cells (MSC), osteoblasts (OB), and high-grade OS biopsies from patients who did (5yr mets) or did not develop metastasis (No mets) within 5 years from diagnoses showing increased *UHRF1* expression in metastasis-prone tumors. (F-G) qPCR analysis of *UHRF1* mRNA in (F) human OS cell lines (143B, SJSA-1, SaOS-2 and U-2 OS) and (G) patient-derived xenografts (PDX1-5). All data are mean ± SD normalized to MSC (n=3). (H-I) Western blot detection of UHRF1 in (H) human OS cell lines, and (I) PDXs tumors, β-actin was used as loading control. ** p < 0.01, *** p < 0.001 by unpaired two-tailed *t* test.

The clinical data led us to examine UHRF1 mRNA and protein levels in 4 different human OS cell lines: 143B, SJSA-1, SaOS-2 (RB-null cell line), and U-2 OS, along with 5 patient-derived orthotopic xenografts (PDX1-5). qPCR analysis of human OS cell lines and PDX showed a significant upregulation of *UHRF1* mRNA levels in all samples except PDX2 when compared to human mesenchymal stem cells (MSC), independent of whether they express RB protein or not (Fig. 2F-G and Supp. Fig. 3A). At the protein level, UHRF1 was highly overexpressed across all analyzed OS subjects compared to MSC controls (Fig. 2H-I).

### *UHRF1* is a direct target of the RB/E2F signaling pathway in the osteogenic lineage

A few studies have reported UHRF1 as a direct E2F1 transcriptional target [28, 29]. Following our observations that UHRF1 is overexpressed in all OS samples analyzed, we aimed to confirm whether UHRF1 is a direct E2F1 target in the osteogenic lineage. We performed chromatin immunoprecipitation (ChIP) analysis in MSC and found E2F1 enrichment at the E2F1 consensus binding motifs located within the UHRF1 promoter (Supp. Fig. 3B). To confirm E2F1 drives *UHRF1* transcription in OS cells, we knocked-down E2F1 using shRNA which resulted in a modest decrease in UHRF1 protein levels in OS cell lines (Supp. Fig. 3C). Similarly, knockdown of another activator E2F, E2F3, showed mild to no effect on UHRF1 protein levels (Supp. Fig. 3D). However, knockdown of E2F1 in combination with E2F3, but not in combination with E2F2, significantly decreased UHRF1 protein levels (Supp. Fig. 3F). To further verify that UHRF1 expression is transcriptionally regulated by the RB/E2F pathway, human OS cell lines were treated with palbociclib. Palbociclib is a CDK4/6 inhibitor that results in blockade of RB hyperphosphorylation and thereby, E2F transcriptional repression. qPCR analysis revealed significant decreases in *UHRF1* mRNA levels upon palbociclib treatment, with the exception of the RB-null cell line, SaOS-2, which served as a negative control (Supp. Fig. 3G). Palbociclib treatment also decreased UHRF1 protein levels in a dose-dependent manner in RB-positive OS cells, as revealed by Western Blot analysis (Supp. Fig. 3H). In line with these results, we also observed upregulation of *CDK4* and/or *CDK6* transcripts in RB-wildtype OS cell lines and/or *INK4A* downregulation (Supp. Fig. 3I-J). SJSA-1 is known to have *CDK4* amplification and U2-OS to have *INK4A* deletion. Together, these data confirm transcriptional regulation of *UHRF1* by the RB/E2F pathway through direct transcriptional activation by activator E2Fs, E2F1 and E2F3.

### UHRF1 promotes human OS proliferation *in vitro* and *in vivo*

Emerging reports have linked UHRF1 overexpression in cancer with its ability to promote proliferation, migration/invasion, or both [11, 30–35]. To study the role of UHRF1 overexpression in OS, we first used shRNA to UHRF1-knockdown in OS cell lines; however, we observed that UHRF1 knockdown could be sustained for more than two weeks, complicating data reproducibility over time (data not shown). To overcome this challenge, we generated UHRF1 CRISPR knockouts (KO) and a non-targeting vector control (VC). Clones (n=3) were selected for SJSA-1 and 143B, while the other cell lines remained as pools and were analyzed within 2-3 weeks after being generated. Western blot analysis revealed successful UHRF1 KO in all four OS cell lines tested (Fig. 3A). Growth curve analyses from these cells showed that UHRF1 KO cells have longer doubling times compared to VC OS cells (Fig. 3B). This decrease in proliferation potential was further evidenced through clonogenic assays showing significant reductions in both the number and the size of colonies in UHRF1 KO compared to VC cells (Fig. 3D). To directly assess cell proliferation, we labelled actively proliferating UHRF1 KO and VC cells using EdU and found significantly fewer EdU-positive UHRF1 KO cells compared to VC, suggesting a slower rate of active DNA synthesis; with the exception of U-2 OS, albeit a trend of decreased incorporation was still observed (Fig. 3E-F).

**Figure 3.**
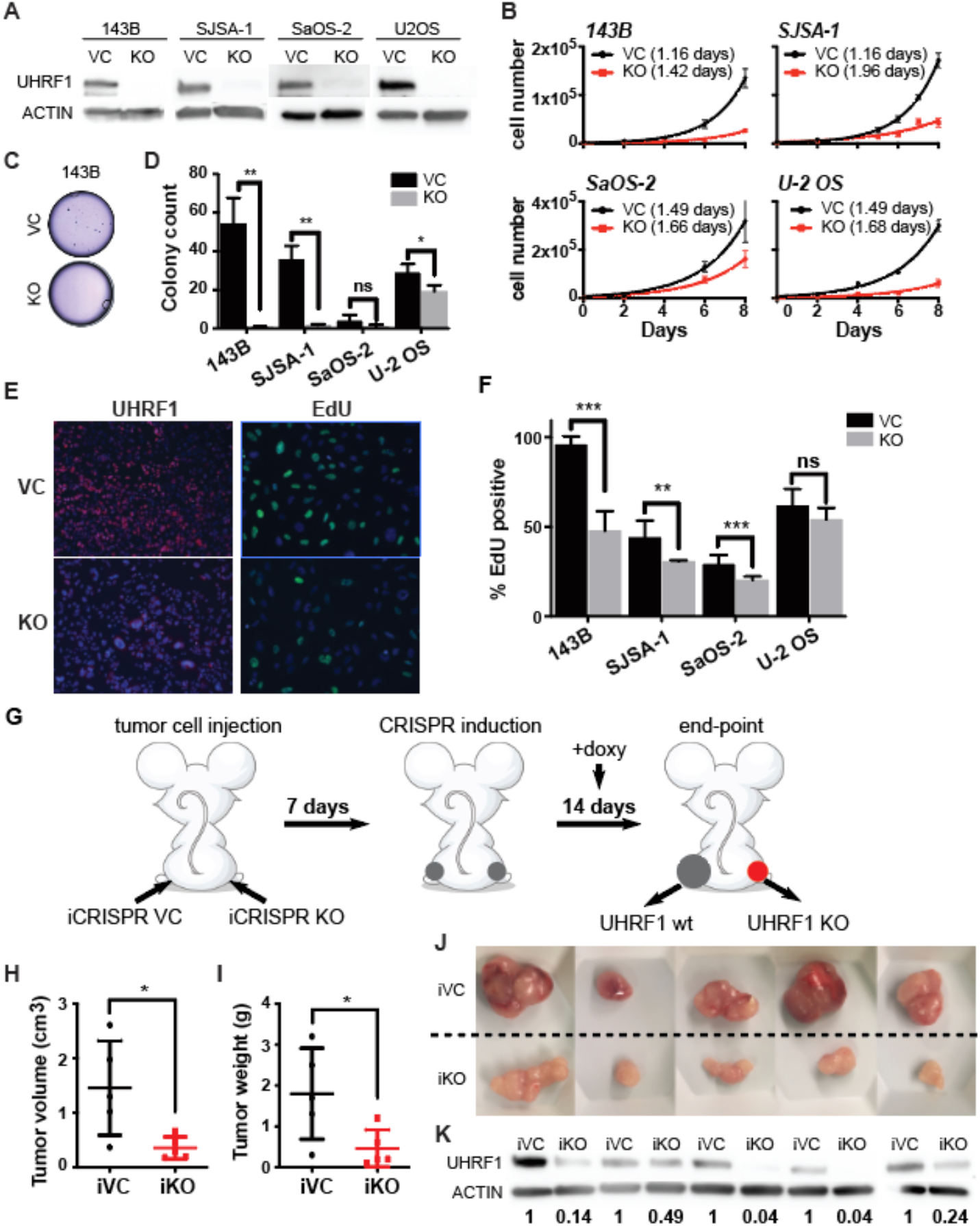
UHRF1 promotes human OS tumor growth *in vitro* and *in vivo*. (A) Western blot verification of CRISPR/Cas9-mediated UHRF1 knockout of human OS cell lines (KO) in comparison to non-targeting vector control (VC). β-actin was used as loading control. (B) Growth curves of UHRF1 knockout cells (KO, red) compared to control (VC, black), population doubling time indicated in parentheses. (C) Representative images from clonogenic assay plates with 143B VC and UHRF1 KO. (D) Histogram of colony counts from clonogenic assay. ns=not significant; * p < 0.05, ** p < 0.01 by unpaired two-tailed *t* test. (E) Representative images of cells labelled with Click-iT EdU (green) or immunostained with UHRF1 (red). Nuclei were counterstained with DAPI (blue). (F) Quantification of EdU signal from immunocytochemistry probing UHRF1 and EdU in VC and UHRF1 KO human OS cell lines. ns=not significant; ** p < 0.01, *** p < 0.001 by unpaired two-tailed *t* test. (G) Cartoon representation for doxycycline-inducible *in vivo* knockout for flank-injected OS cell lines carrying a non-targeting sgRNA (iCRISPR VC) or UHRF1 sgRNA (iCRISPR KO). (H-I) Quantification of the tumor (H) volume and (I) weight for each of the replicates. * p<0.05 by paired two-tailed *t* test. (J) Images from tumors collected from subcutaneous injection of iCRISPR VC (iVC, n=5) and iCRISPR KO (iKO, n=5) SJSA-1. (K) Western blot verification of UHRF1 knockout in iKO compared to iVC for tumors shown in J. β-actin was used as loading control. UHRF1 densitometry quantification normalized to the matching iVC, prior normalization to β-actin.

To assess proliferative capacity *in vivo*, SJSA-1 UHRF1 KO and VC OS cells were injected subcutaneously into immune-deficient mice at opposite flank regions. UHRF1 KO cells gave rise to significantly smaller tumors (average size 0.49 ± 0.37 cm^3^ and average weight 0.70 ± 0.69 g) when compared to VC (average volume 1.23 ± 0.75 cm^3^ (p=0.02, n=10); and average weight 1.62 ± 1.09 g (p=0.01, n=10)). To test the therapeutic potential of targeting UHRF1 in established tumors, we utilized a doxycycline-inducible system to drive the expression of CRISPR gRNA against UHRF1. Inducible UHRF1 KO (iKO) and VC (iVC) OS cells were subcutaneously injected, and tumors allowed to establish for one week. CRISPR/Cas-9 gRNAs were then induced using oral doxycycline (Fig. 3G). SJSA-1 tumor growth was significantly reduced in UHRF1 iKO tumors (average tumor volume 0.356 ± 0.201 cm^3^ and weight 0.46 ± 0.45 g) compared to iVC tumors (1.455 ± 0.865 cm^3^ (p=0.02, n=5); and 1.80 ± 1.11 g (p=0.01, n=5)) (Fig. 3H-I). Western blot analyses confirmed reduced UHRF1 protein levels in UHRF1 iKO tumors (Fig. 3K). UHRF1 iKO tumors also presented phenotypic features of less aggressive tumors, as seen by lessened vasculature and ulceration compared to iVC tumors (Fig. 3J). Induction of a different UHRF1 sgRNA in SJSA-1 resulted in similar tumor size reduction, as well as with other OS cell lines (Supp. Fig. 4 A-D). Taken together, both *in vitro* and *in vivo* data suggest that UHRF1 is critical for OS cell proliferation.

### UHRF1 promotes human OS cell migration and invasion *in vitro* and *in vivo*

The high metastatic potential of OS is the main cause of mortality in patients. Given that our mouse models indicated a possible involvement of UHRF1 in OS metastasis, we sought to evaluate the role of UHRF1 in promoting cell migration, invasion, and metastasis. Scratch-wound assays were used to assess the migratory potential of UHRF1 KO OS cell lines (Fig. 4A). We observed a significant decrease in the distance migrated in UHRF1 KO compared to VC OS cells for all cell lines studied, with an average of 56.8 ± 2.3% reduction in migration across the three cell lines (Fig. 4B). In addition to decreased migratory potential, Transwell invasion assays revealed a significant reduction in the number of UHRF1 KO OS cells capable of mobilizing across Matrigel-coated inserts in all OS cell lines (Fig. 4C-D).

**Figure 4.**
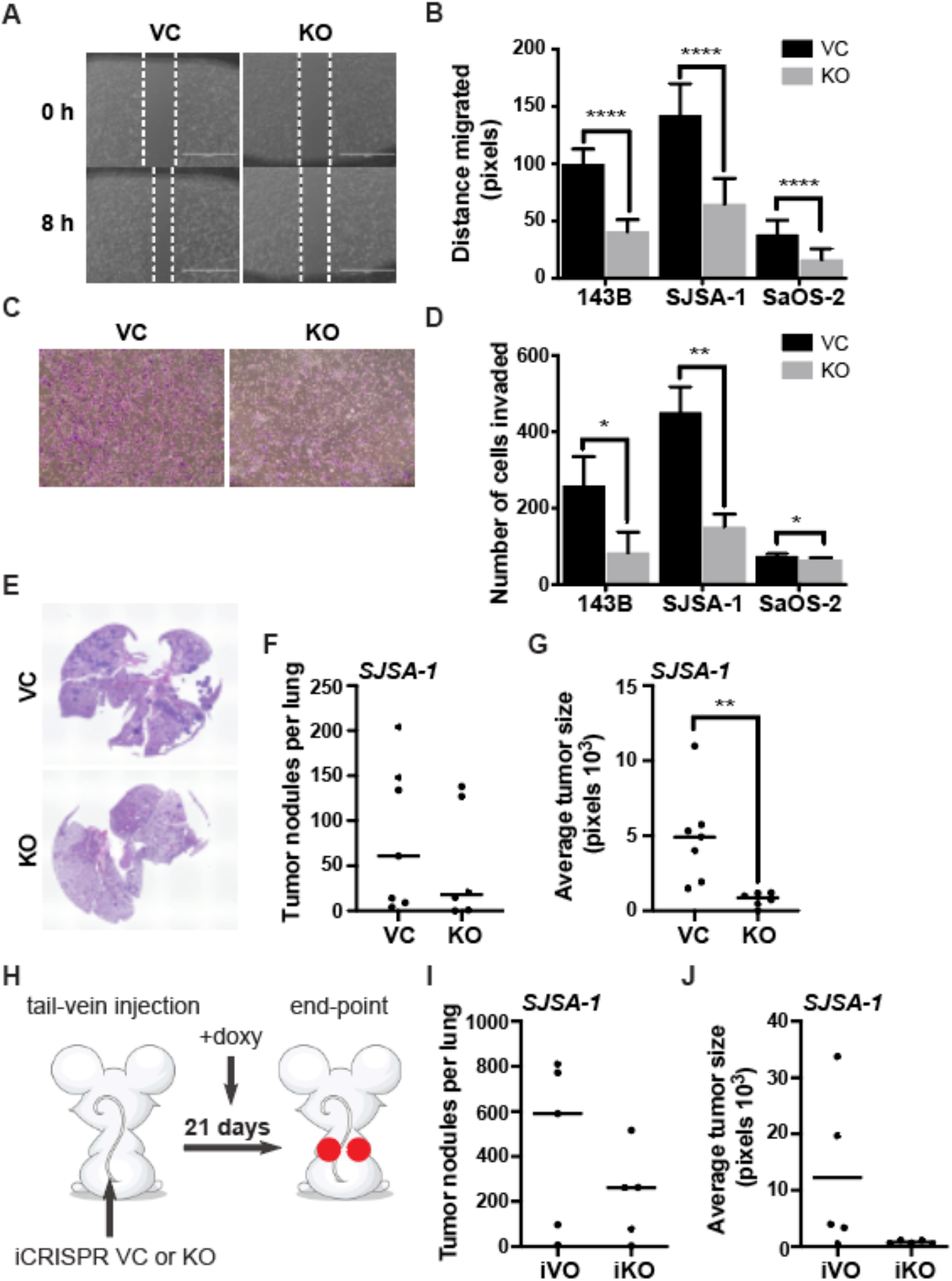
UHRF1 promotes human OS cell migration and invasion *in vitro* and *in vivo*. (A) Representative image from scratch-wound assay comparing wound closure of non-targeting vector control (VC) and UHRF1 KO (KO) cells over 8 h in SJSA-1 cells. White dashed lines represent wound edge. (B) Quantification of distance (pixels) migrated in scratch-wound assays for each of the OS cell lines. Each data point is mean ± s.d. of ten measurements in triplicate samples. (C) Representative images from Transwell assays with cells stained with crystal violet (purple) comparing levels of invasion between VC and UHRF1 KO in SJSA-1 cells. (D) Quantification of number of cells invaded across the Transwell membrane for each of the OS cell lines. Each data point is mean ± s.d. of triplicate samples. (E) Representative H&E staining from lung sections collected from mice injected intravenously with VC or UHRF1 KO SJSA-1 cells. Pulmonary nodules were quantified by (F) the number of tumor nodules found per lung and by (G) the average tumor size per lung. Solid line represents the average from n=7 for VC and n=6 for KO. (H) Cartoon representation for doxycycline-inducible *in vivo* knockout for tail-vein-injected OS cell lines carrying a non-targeting sgRNA (iCRISPR VC; iVC) or UHRF1 sgRNA (iCRISPR KO; iKO) to assess lung metastases. (I-J) Pulmonary nodules were quantified by (I) the number of tumor nodules found per lung and by (J) the average tumor size per lung. Solid line represents the average from five replicates for each condition. For all graphs: ** p < 0.01, *** p < 0.001, **** p < 0.0001 by unpaired two-tailed *t* test.

To test whether the reduced migratory and invasive capacities observed *in vitro* translate into less metastatic potential *in vivo*, we injected SJSA-1 UHRF1 KO and VC cells into the tail vein of immune-deficient mice and assessed the rate of lung colonization 3-weeks post injection. We found that the lungs of mice injected with SJSA-1 UHRF1 KO cells had a reduced number of metastatic nodules and a significantly reduced tumor nodule size when compared to the lungs of mice injected with VC cells (Fig. 4E-G). We also observed a significant decrease in the burden of lung metastases when we injected SJSA-1 UHRF1 iKO into the tail-vein of mice fed with doxycycline, compared to iVC cells (Fig. 4H-J), as reflected by the reduced number (Fig. 4I) and significant decrease in size (Fig. 4J) of metastatic nodules. This suggests that targeting UHRF1 would be an effective strategy to reduce the burden of lung metastases even when tumor cells have already undergone intravasation.

To determine whether UHRF1 plays an intrinsic role in cell migration, we performed gain-of-function studies using MSCs, the normal progenitors of the osteogenic tissue. We generated a doxycycline-inducible lentiviral vector system to drive non-toxic UHRF1 overexpression (pCW57-UHRF1; Fig. 5A) and used the empty vector as control (pCW57-VC). Western blot analysis confirmed successful induction of UHRF1 overexpression upon doxycycline treatment in transduced MSCs (Fig. 5B). We found that UHRF1 overexpression in MSCs significantly increased migratory potential as evidenced by the robust increase in the distance migrated (Fig. 5C-D) and number of cells invaded across Matrigel-coated Transwell membranes (Fig. 5E-F) upon doxycycline induction in pCW57-UHRF1 cells compared to pCW57-VC using scratch-wound assay.

**Figure 5.**
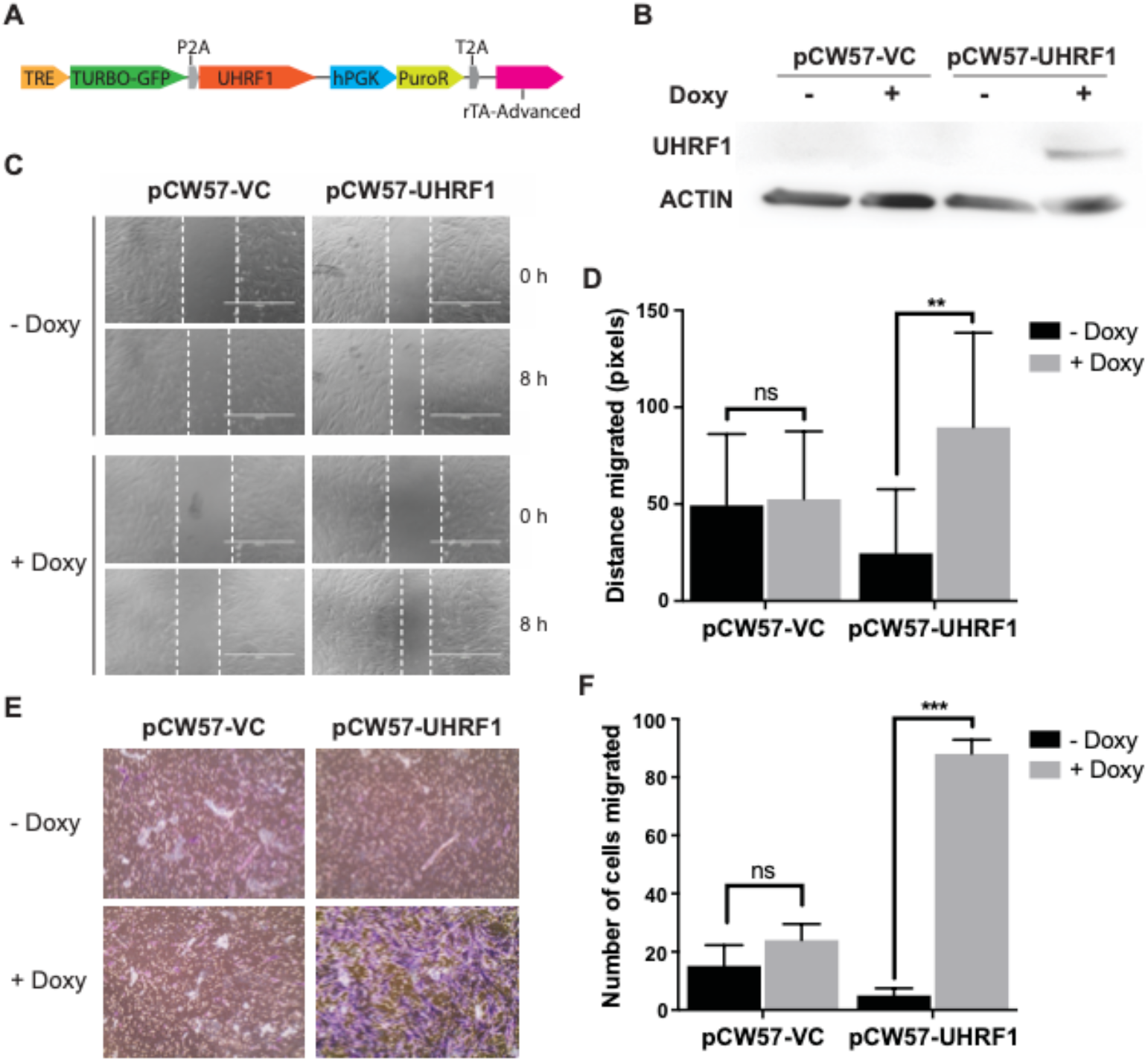
UHRF1 overexpression promotes human MSC cell migration and invasion. (A) Plasmid map of the doxycycline inducible UHRF1 overexpression system. Tet Response Element (TRE) is activated by doxycycline binding to the reverse tetracycline-controlled transactivator (rTA), allowing transcription of GFP and the pCW57-promoter driven human UHRF1 cDNA. (B) Western blot detection of UHRF1 in MSCs transduced with vector control (pCW57-VC) or UHRF1 overexpression (pCW57-UHRF1) plasmid, treated with DMSO (-) or doxycycline (+). (C) Scratch-wound assay in MSC culture comparing wound closure of control (pCW57-VC) and UHRF1 overexpressed (pCW57-UHRF1) cells over 8 h span. White dashed lines represent wound edge. (D) Histogram quantification of distance (pixels) migrated for doxycycline non-induced (black) and induced (gray) pCW57-VC and pCW57-UHRF1. Each data point is mean ± s.d. of ten measurements in triplicate samples. (E) Representative images from Transwell assays with cells stained with crystal violet comparing levels of invasion between doxycycline non-induced and induced pCW57-VC and pCW57-UHRF1 MSC. (F) Quantification of the number of cells invaded across the Transwell membrane. Each data point is mean ± s.d. of triplicate samples. For all graphs: ns=not significant, ** p < 0.01, *** p < 0.001, by unpaired two-tailed *t* test.

We utilized this pCW57-UHRF1 vector to rescue high UHRF1 expression in UHRF1 KO OS cells. Doxycycline treatment for 24 h was able to restore UHRF1 protein levels back to wild-type levels (Supp. Fig. 4E). The rescue of UHRF1 expression resulted in a significant increase in migration in all OS cell lines (Supp. Fig. 4F). Taken together, this results support a role of UHRF1 in OS migration and invasion.

### UHRF1 increases exosome and *PLAU* expression resulting in enhanced cell migration and invasion

UHRF1 is required for maintenance of DNA methylation during DNA replication through the recruitment of DNMT1 [36]. However, UHRF1 ubiquitin ligase activity also targets DNMT1 for degradation, with some studies reporting that UHRF1 overexpression results in decreased DNA methylation [37, 38]. To determine the effects of UHRF1 loss on DNA methylation, chromatin structure, and transcription, we analyzed global DNA methylation, chromatin accessibility, and transcriptome changes. We probed genomic DNA isolated from UHRF1 KO and VC cell lines with an antibody against 5-methyl cytosine (5meC) to assess global DNA methylation. We found genomic 5meC levels were decreased in UHRF1 KO compared to VC across all OS cell lines examined (Supp. Fig. 5A). This reduction in global DNA methylation in OS UHRF1 KO cells compared to VC was confirmed by representation bisulfite sequencing analysis of differentially methylated regions across the genome of VC and UHRF1 KO OS cells, which showed most of the changes occurring at non-coding regions of the genome (Supp. Fig. 5B). We also analyzed changes in the chromatin landscape upon loss of UHRF1 using ATAC-seq in 143B, SJSA-1, and SaOS-2 UHRF1 KO compared to VC (Supp. Fig. 5C). Despite the role of UHRF1 in heterochromatin maintenance and the decreased genomic levels in DNA methylation, we identified only 16 chromatin regions with significant changes in chromatin accessibility across these 3 biological replicates (Supp. Table 2). Within these changes, the majority were consistent with the role of UHRF1 chromatin repression, with approximately 69% (11/16) resulting in opening of chromatin upon UHRF1 loss.

In line with modest changes in DNA methylation and chromatin accessibility, gene expression analysis using RNA-seq revealed that the gene expression profile from UHRF1 KO cell lines present minor variability from VC cells, as portrayed by the principal component analysis (PCA) plot (Supp. Fig. 5D). We identified only 272 differentially expressed genes (DEGs), with 191 upregulated and 81 downregulated genes in UHRF1 KO compared to VC cells. Of particular interest, gene ontology (GO) analysis for biological processes of the downregulated DEGs showed an enrichment of genes involved in regulation of cell adhesion and migration (Fig. 6A). GO analysis of cellular components exposed the enrichment of genes involved in extracellular vesicle formation and cell adhesion, consistent with the less migratory phenotype of UHRF1 KO OS cells (Fig. 6B). We confirmed the changes in gene expression observed among top 13 genes involved in regulation of cell migration (Fig. 6C) through qPCR (Fig. 6D).

**Figure 6.**
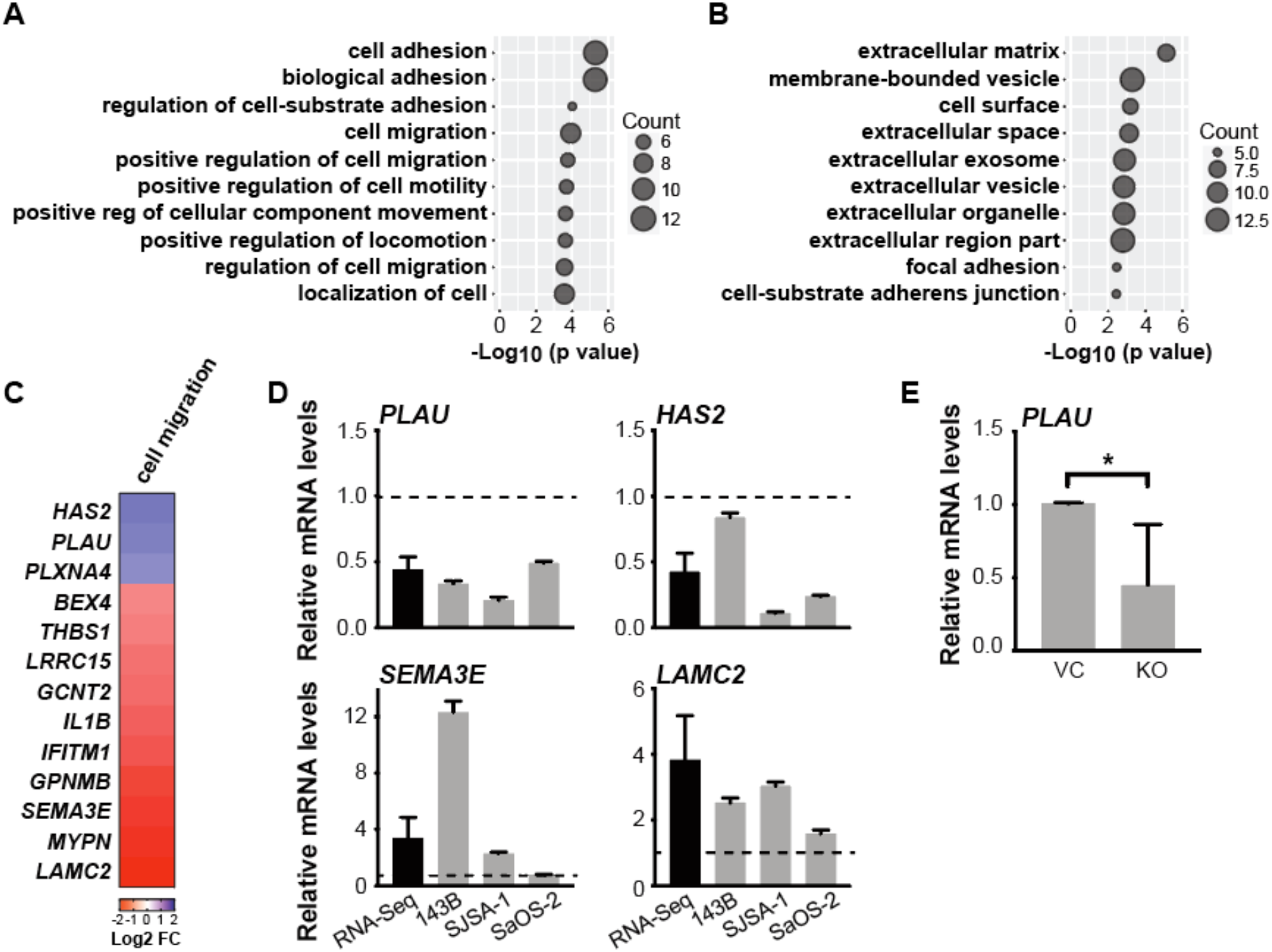
UHRF1 controls the expression of genes involved in exosomes and cell migration and invasion. (A) Gene ontology (GO) analysis for biological processes and (B) cellular components of the downregulated differentially expressed genes (DEGs) identified through RNA-seq. (C) Heat map of top 13 genes involved in regulation of cell migration, decreased (blue) or increased (red) in expression level upon UHRF1 loss. (D) qPCR analysis confirming expression changes of migration related genes in individual cell lines upon UHRF1 loss, fold change of mRNA level normalized to vector control (gray), compared with average fold change from RNA-seq analysis (black). (E) qPCR analysis of *PLAU* mRNA levels in flank-implanted tumors from control (VC) and UHRF1 KO (KO) SJSA-1 cell lines. * p<0.05 by paired two-tailed *t* test.

To examine whether UHRF1 affects migration and invasion through exosomes, we first tested the differential effects of conditioned media (24 h) from VC and UHRF1 KO cells on OS cell migration. We found that conditioned media from OS VC cells significantly increased autologous OS cell migration compared to fresh media (Fig. 7A, blue bars). While conditioned media derived from OS UHRF1 KO cells also increased cell migration compared to fresh media, the VC cell migration when exposed to UHRF1 KO conditioned media was lower than VC conditioned media (Fig. 7A, red bars). Interestingly, UHRF1 KO cells also displayed a robust increase in cell migration when exposed to VC conditioned media, but not when exposed with autologous UHRF1 KO conditioned media (Fig. 7B). This suggests that UHRF1 loss decreases the secretion of pro-migratory factors to the extracellular matrix that can serve as an autologous signal to stimulate OS cell migration but does not affect the ability of the cells to respond to these extracellular factors. We next blocked exosome biogenesis/release using GW4869 and endocytosis using chlorpromazine (CPZ). We found that both GW4869 and CPZ significantly reduced the cell migration induced by OS conditioned media (Fig. 4C). Taken together, this indicates that UHRF1 overexpression contributes to the expression and secretion of exosomes which contribute, at least in part, to increased OS cell migration.

**Figure 7.**
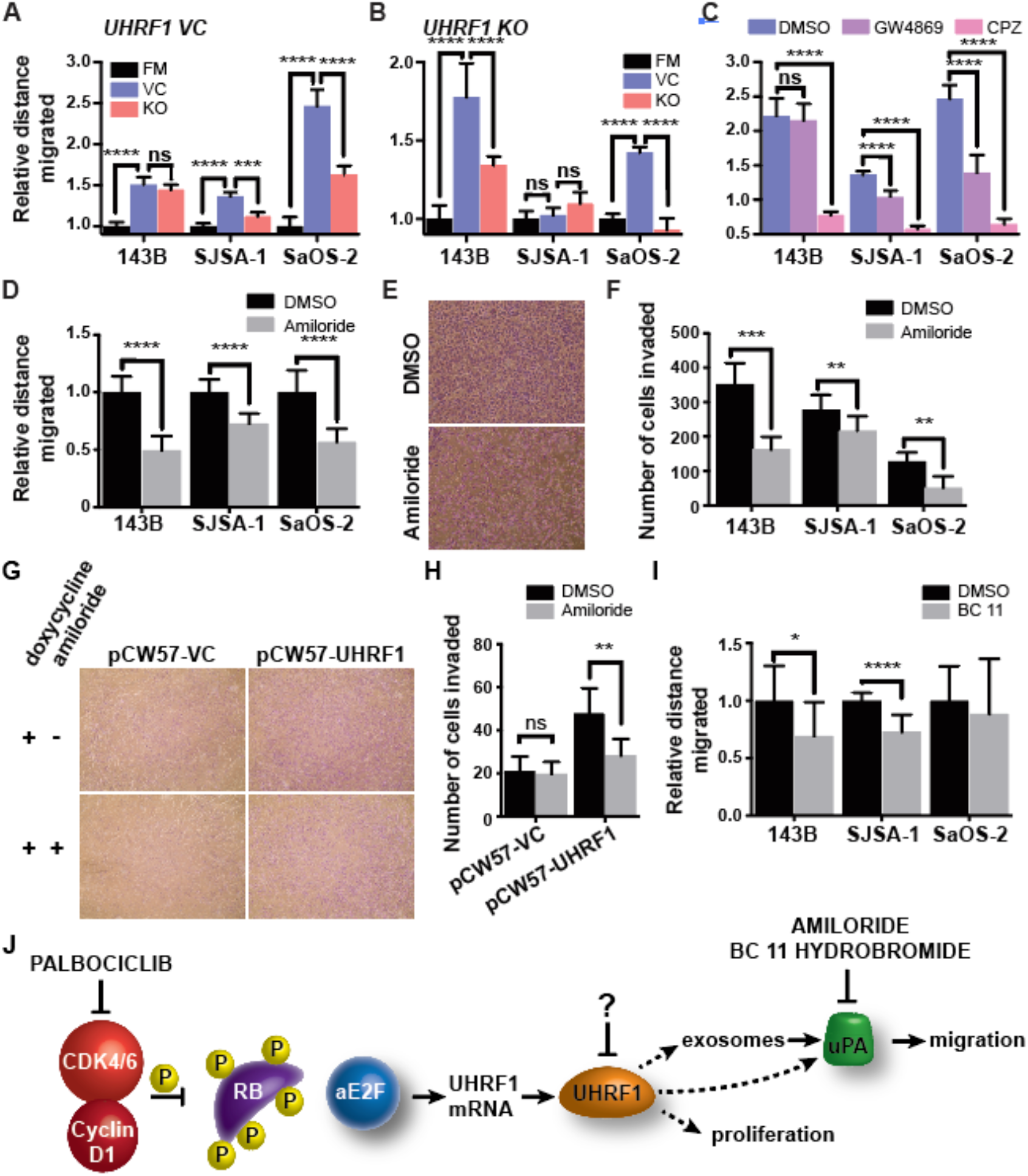
Exosome release and urokinase plasminogen activator contribute to OS migration. (A-D) Quantification of relative distance migrated in scratch-wound assays for (A) OS VC cells assayed with fresh media (FM) or conditioned media from VC cells or UHRF1 KO cells, normalized to FM; (B) OS UHRF1 KO cells assayed with FM or conditioned media from VC cells or UHRF1 KO cells, normalized to FM; (C) each of the OS cell lines treated with DMSO, 10 μM GW4869 or 20 μM CPZ, normalized to DMSO control; (D) each of the OS cell lines treated with DMSO or 150 μM amiloride, normalized to DMSO control. Each data point is mean ± s.d. of ten measurements in triplicate samples. (E) Representative images from Transwell assays with cells stained with crystal violet comparing levels of invasion between cells treated with DMSO or 150 μM amiloride in SJSA-1. (F) Quantification of the number of cells invaded in Transwell invasion assay for each of the OS cell lines treated with DMSO or 150 μM amiloride. Each data point is mean ± s.d. of triplicate samples. (G) Representative images from Transwell assays with cells stained with crystal violet comparing levels of invasion between doxycycline induced pCW57-VC and pCW57-UHRF1 MSC treated with DMSO or 150 μM amiloride. (H) Quantification of the number of MSC invaded across the Transwell membrane from experiment shown in G. Each data point is mean ± s.d. of triplicate samples. (I) Quantification of distance (pixels) migrated in scratch-wound assays for each of the OS cell lines treated with DMSO or 16.4 μM BC 11 hydrobromide in SJSA-1, normalized to DMSO control. Each data point is mean ± s.d. of ten measurements in triplicate samples. For all graphs: ns=not significant, ** p < 0.01, *** p < 0.001, by unpaired two-tailed *t* test. (J) Model of UHRF1 oncogenic function in OS. In the absence of RB or upon RB inactivation via hyperphosphorylation, activator E2Fs drive *UHRF1* transcription. The resulting UHRF1 protein overexpression stimulates proliferation and uPA overexpression, aiding to migration, invasion, and metastasis. For OS with CDK4/6 amplification, palbociclib treatment is an effective way to reduce UHRF1 expression and result in decreased proliferation and migration. Downstream of UHRF1, uPA inhibitors (e.g. amiloride, BC 11 hydrobromide) are attractive therapeutic options to decrease migration and metastasis. For most OS (with or without *RB1* mutations), the development of UHRF1-targeted therapeutics might result in beneficial decrease in both tumor growth and pulmonary metastases.

Plasminogen activator, Urokinase (*PLAU*) was one of the genes identified through the transcriptome analysis to be potentially involved in UHRF1-mediated cell migration. Urokinase plasminogen activator protein (uPA) is associated with cell migration and a known cargo protein in osteosarcoma-secreted exosomes [39]. Significant decrease in the levels of the *PLAU* transcript were observed in UHRF1 KO OS cells (Fig. 6D), as well as the tumors they form subcutaneously in immune-deficient mice, with an average 2.2-fold decrease when compared to tumors formed from VC cells (p=0.038, Fig. 6E). Thus, we tested whether treatment with uPA inhibitors could decrease migratory potential and invasiveness of OS cells. Scratch-wound assays of amiloride-treated OS cell lines showed a significant decrease in migratory potential, with an average of 40.3 ± 11.9% decrease across all OS cell lines examined (Fig. 7A). A significant decrease of invasiveness, as determined by Transwell invasion assay, was also observed upon amiloride treatment (Fig. 7B-C). This confirmed uPA as an important contributor of OS migratory potential. In line with our observations associating UHRF1 with uPA levels, amiloride treatment was also able to inhibit UHRF1-induced cell migration in MSCs (Fig. 7D-E). BC 11 hydrobromide, a selective uPA inhibitor, was used to further verify the role of uPA in OS cell migration. A 23.2 ± 10.2% decrease in migration is seen across OS cell lines upon treatment (Fig. 7F). Taken together, we established a positive correlation between UHRF1 expression levels and migration/invasion potential in the osteogenic lineage, with uPA as the potential conduit between the two.

## DISCUSSION

Metastasis remains the most frequent fatal complication of OS. The lack of significant breakthroughs beyond the scope of standard chemotherapeutics targeting highly proliferative underscores a pressing clinical need for new therapeutic strategies for OS. For decades, genetic alterations at the *RB1* gene have been associated with increased mortality, metastasis and poor response to chemotherapy in OS [6–10, 40, 41]. However, the precise mechanism through which this occurs remains to be elucidated to facilitate the development of better therapeutic strategies. In this study, we identified *UHRF1* as a direct target of RB/E2F pathway. UHRF1 overexpression is directly correlated with increased OS malignancy and metastatic disease in humans. Specifically, UHRF1 facilitates OS cell proliferative, invasive, and metastatic capacity. Furthermore, we found that the decrease in survival associated with loss-of-function mutations at the *Rb1* gene is critically mediated through the overexpression of UHRF1.

Studies from retinoblastoma, a cancer initiated by biallelic inactivation of the *RB1* gene, defined a central role of RB in epigenetic control and a key role of RB/E2F-regulated chromatin remodelers in tumorigenesis [11, 42, 43]. Among these chromatin remodelers, UHRF1 was identified as a potential critical regulator of tumor initiation and progression following RB-pathway deregulation that is overexpressed in several cancers [11–15]. We confirmed *UHRF1* upregulation in human OS cell lines, PDXs, and primary OS tissue. UHRF1 protein overexpression is recapitulated in both *p53* cKO and *Tp53*/*Rb1* DKO OS mouse model, validating the suitability of this mouse model for the study of OS. Direct regulation of UHRF1 by the RB/E2F signaling pathway was described previously, though in limited context [28, 29]. We established that *UHRF1* expression is transcriptionally regulated by the RB/E2F signaling pathway in the osteogenic lineage. ChIP analysis revealed enrichment of E2F1 at putative binding motifs within the promoter region of *UHRF1*. However, E2F1 knockdown was insufficient in reducing UHRF1 expression. Rather, significant reduction of UHRF1 level was achieved only through combined knockdown of E2F1 and E2F3, suggesting compensatory mechanisms between activator E2Fs in regulating the transcription of downstream targets. As the RB/E2F signaling pathway is often inactivated in cancer through mechanisms other than *RB1* loss-of-function mutations, the most common being RB hyperphosphorylation through CDK4/6 activation or p16 inactivation [17, 44, 45], it is not surprising that UHRF1 upregulation and overexpression in OS is not restricted to *RB1*-null cases. UHRF1 overexpression in *RB1*-wildtype OS cell lines results from the upregulation of *CDK4* and/or *CDK6*, which leads to RB hyperphosphorylation, preventing the repression of E2F binding activities. This was confirmed by the down-regulation of *UHRF1* and dose-dependent decrease of UHRF1 upon treatment with palbociclib, a known CDK4/6 inhibitor [46].

UHRF1 expression is positively correlated with cell proliferation and cell mobility in both normal and malignant cells *in vitro* [11, 32–35]. Although it is clear that UHRF1 expression is linked to cellular phenotypes that strongly associate with tumor malignancy, the link between UHRF1, OS tumor progression, and its role in the poor prognosis associated with *RB1* loss has not been established. Our results showed that UHRF1 KO decreases clonogenicity and tumor growth both *in vitro* and *in vivo*. Independent from its role on proliferation, UHRF1 has a clear effect on the migratory potential of the osteoblastic lineage as UHRF1 overexpression was sufficient to increase migration and invasion of normal MSCs and targeting UHRF1 drastically reduced migration, invasion and metastatic capacity of OS cell lines. It is important to note that although *RB1*-null status has been clinically associated with poor disease outlook, the *RB1*-null cell line, SaOS-2, does not exhibit higher aggressiveness in comparison to other cell lines. This could be explained by the limitation of utilizing cell lines, in which loss of clinical representation is commonly seen. Our results also present a novel role of UHRF1 in the control of exosome formation and suggest that uPA is critical for UHRF1-mediated migration. Exosomes and their cargo have been implicated in osteosarcoma metastasis [47]. Here we show that UHRF1 can alter the expression of exosomal components as well as their protein cargo composition. uPA is a serine protease that has long been associated with tumor malignancy, being associated with cell migration and a known cargo protein in osteosarcoma-secreted exosomes [39]. Upon conversion of plasminogen into plasmin, uPA triggers a proteolytic cascade that leads to the degradation of extracellular matrix, an event necessary for angiogenesis and metastasis. Our findings are in line with a previous study describing the uPA/uPAR axis as a metastatic driver in OS [48]. While our results suggest that targeting UHRF1 might result in a more robust therapeutic outcome, the observation that the UHRF1 induced cell migration can be reversed by treatment with uPA inhibitors like amiloride [49] and BC11 hydrobromide, provides a therapeutic alternative that can be further explored.

Our group is the first to model the critical role of UHRF1 in osteosarcomagenesis, with *Uhrf1* KO resulting in a dramatic increase in overall survival and decrease in metastases, particularly in *Rb1*-null OS. Indeed, median survival and rate of metastasis of *Tp53*/*Rb1*/*Uhrf1* TKO closely resembled that of *Tp53* cKO mice, indicating that UHRF1 is a major driver of *Rb1* loss-associated malignancy. Interestingly, our results also indicate that the increased survival in *Tp53*/*Rb1*/*Uhrf1* TKO mice is largely a result of delayed tumor promotion, with negligible differences in tumor growth rates once the tumors have been detected. This is quite distinct from our observations targeting UHRF1 in established tumor cells, suggesting there might be compensatory pathways arising in the developmental tumor models. Further studies on tumors arising from these developmental OS models might provide insight on potential compensatory mechanisms of *Uhrf1* loss which may be relevant for predicting mechanisms of resistance for future UHRF1-targeted therapeutics.

While targeting *Uhrf1* in developmental mouse models showed a distinct improvement in survival in *Rb1*-null tumors over *Rb1*-wildtype, such distinctions were not observed in UHRF1 overexpressing human OS cell lines. Both *RB1*-null and *RB1*-wildtype OS cell lines appear to equally benefit from targeting UHRF1. It is possible that the timing of UHRF1 overexpression is crucial for the resulting poor prognosis. Early *Rb1* loss would result in immediate UHRF1 overexpression which quickly promotes tumorigenesis compared to tumors that eventually inactivate the RB/E2F pathway through alternative mechanisms. Supporting this theory, *Tp53/Uhrf1* DKO mice showed some improvement in survival compared to the more aggressive *Tp53* cKO mice that showed RB/E2F pathway inactivation. Alternatively, the RB/E2F complex is known not only for its role in direct transcriptional repression, but also as epigenetic modifiers through the recruitment of chromatin remodelers [42, 50–53]. Two UHRF1 domains, PHD and RING, contain LXCXE RB-binding motifs [54]. Unpublished data from our group shows UHRF1 preferentially binds to hyperphosphorylated RB. Thus, *RB1*-wildtype OS cells may preserve protein-protein interactions and RB regulation on UHRF1. The possible loss of this layer of regulation in *RB1-*null tumors could account for the drastic difference in clinical prognosis, supporting the view that the inactivation of RB by phosphorylation is not functionally equivalent to the mutation of the *RB1* gene [55]. Further unpublished data from our group strongly supports a critical role of UHRF1 in aiding RB-mediated tumorigenesis, with loss of *Uhrf1* completely abrogating tumor formation in an *Rb1/p107* knockout model of retinoblastoma and significantly increasing survival of a *Tp53/Rb1* model of small cell lung cancer, similar to the observations presented for OS in this study.

In summary, this study provides evidence supporting UHRF1 targeting as a novel therapeutic strategy in treating OS. We showed that UHRF1 targeting holds great potential in overcoming the highly metastatic characteristic of OS, the main reason why survival rate has remained stagnant in past decades. We stand to be the first to model the effect of UHRF1 throughout development and showed minimal on-target toxicity, as opposed to various chemotherapeutic agents known to affect bone homeostasis [56]. Moreover, the degree to which UHRF1 loss reverted the increased malignancy upon *Rb1* loss suggests that the benefit of UHRF1 targeting could go beyond the scope of OS by improving current treatment paradigms of other cancers harboring RB/E2F pathway inactivation.

## ACKNOWLEDGEMENTS

We thank Erin Yamanaka, Josselyn Peña, Sara Akhlaghi, and Annie Jeon for help with mouse genotyping and cell culture; Christopher Liang for help with FDG-PET/microCT scans; Dr. Ali Nael, Dr. Robert Edwards and Dr. Aaron Sassoon for assistance with pathology analysis. This work was supported in part by grants to C.A.B. from the NIH (CA178207 and CA229696), the American Cancer Society (129801-IRG-16-187-13 and 133403-RSG-19-031-01-DMC) and the 2018 AACR-Aflac, Inc. Career Development Award for Pediatric Cancer Research (18-20-10-BENA). J.L. and K.S. were supported by the NIH-MARC U-STAR training grant (T34GM136498). This work was also made possible, in part, through access to the Genomics High Throughput Facility Shared Resource of the Cancer Center Support Grant (P30CA-062203) at the University of California, Irvine and NIH shared instrumentation grants 1S10RR025496-01, 1S10OD010794-01, and 1S10OD021718-01.

## Author contributions

C.A.B. conceived the project. S.C.W., A.K., L.Z., J.L., K.S., and C.A.B. performed the experiments. J.W. and C.C. performed the sequencing data analysis. S.W. and C.A.B. wrote the manuscript.

