## Supplemental Data for "UHRF1 upregulation mediates exosome release and tumor progression in osteosarcoma"

**Supplemental Figure Legends**

**Supplemental Figure 1.** (A) FDG-PET/microCT scans from the three *Tp53/Rb1/Uhrf1* TKO mice that had not acquired tumors well beyond the average of the rest of their study group (> 69 weeks) show no signs of detectable tumors. (B-F) qPCR analysis of (B) *Uhrf1*, (C) Cdk4, (D) *p16*, (E) *Cdk6*, and (F) *RB1* mRNA expression in osteosarcoma mouse tumors from the 70% lower age to end-of-study (young; n=8) compared to the older 30% (old; n=3) mice. Data are mean ± SD normalized to mouse MSC.

**Supplemental Figure 2.** Representative H&E staining from OS tumor sections collected from genetically engineered osteosarcoma mouse models subjected to pathology analysis: *Tp53/Rb1* DKO: *Osx-Cre;* $p53^{lox/lox}$; $Rb1^{lox/lox}$ , *Tp53/Rb1/Uhrf1* TKO: *Osx-Cre;* $p53^{lox/lox}$; $Rb1^{lox/lox}$ ;$Uhrf1^{lox/lox}$ , *Tp53* cKO: *Osx-Cre;* $p53^{lox/lox}$; $Rb1^{lox/lox}$ and *Tp53/Uhrf1* DKO: *Osx-Cre;* $p53^{lox/lox}$; $Uhrf1^{lox/lox}$ . Scale bar = 100 µm.

**Supplemental Figure 3.** (A) Representative Western blot analysis of RB protein presence in human OS cell lines and PDXs. β-actin was used as loading control. (B) Chromatin immunoprecipitation (ChIP) assay reveals enrichment of E2F1 at the three putative binding motifs (M1-3) within *UHRF1* promoter, as predicted by MotifMap. (C-F) Western blot analysis of UHRF1 protein in OS cells transduced with lentivirus containing shRNA against (C) E2F1 (shE2F1), (D) E2F3 (shE2F3), (E) E2F1 and E2F2 (shE2F1/2), and (F) E2F1 and E2F3 (shE2F1/3) compared to scrambled control (shScrmbl). β-actin was used as loading control. Band intensities were quantified by densitometry and normalized to scrambled control. (G) qPCR analysis of *UHRF1* mRNA expression using increasing doses of palbociclib. Data are mean ± SD normalized to DMSO control (n=3). (H) Western blot analysis of UHRF1 protein level in OS cells treated with increasing doses of palbociclib. (I-J) qPCR analysis of (I) *CDK4* and *CDK6* and (J) *INK4A* mRNA expression. Data are mean ± SD normalized to MSC1. *$p < 0.05$, ** $p < 0.01$, *** $p < 0.001$ by unpaired two-tailed *t* test.

**Supplemental Figure 4.** (A-B) Quantification of the SJSA-1 iCRISPR VC (iVC, n=5) and iCRISPR KO gRNA2 (iKO, n=5) tumor volume over time (A) and final tumor volume (B) for each of the replicates. (C-D) Quantification of the 143B iCRISPR VC (iVC, n=5) and iCRISPR KO (iKO, n=5) tumor volume over time (C) and final tumor volume (D) for each of the replicates. Data are mean ± SD. *p < 0.05, ** p < 0.01, *** p < 0.001 by paired two-tailed *t* test. (E) Western blot detection of UHRF1 in OS wild-type and OS UHRF1 KO cells transduced with the inducible UHRF1 overexpression (pCW57-UHRF1) plasmid, treated with DMSO (-) or doxycycline (+). Relative quantification of the UHRF1 levels are shown in the bottom, normalized to wild-type levels. (F) Histogram quantification of distance (pixels) migrated for doxycycline non-induced (black) and induced (gray) pCW57-UHRF1. Each data point is mean ± s.d. of ten measurements in triplicate samples. *p < 0.05, ** p < 0.01, by unpaired two-tailed *t* test.

**Supplemental Figure 5.** (A) Dot blot detection of 5-methyl cytosine signal in non-targeting vector control (VC) and UHRF1 KO (KO) OS cells, comparing global methylation levels. (B) Heatmap for differentially methylated regions generated from RRBS across the genome of VC and KO OS cells. (C) Chromatin landscape generated from ATAC-seq showing peaks of chromatin accessibility across the genome of VC and KO OS cells.

### Supplemental Materials and Methods

#### Chromatin immunoprecipitation

ChIP assays were performed as previously described [17]. ChIP DNA was analyzed by qPCR with SYBR Green (Bio-Rad) in ABI-7500 (Applied Biosystems) using the following primers: Forward: 5'-

CACCCTCTTTCTCGCTTCC-3′; Reverse 5′-TGCCAGCTGCTCTGATTT-3′ (spanning M1); Forward: 5′-CCACATTCCCTCGCAGTATTTA-3′; Reverse 5′-CCCTGAACTCTTAAGTCCAAGTC-3′ (in close proximity with M2, 3). The antibodies used were anti-E2F1 (3742; Cell Signaling) and rabbit IgG (sc-2027, Santa Cruz Biotechnologies).

**5mC DNA dot blot**

Genomic DNA was extracted from cells (Wizard SV genomic DNA purification Kit). Purified DNA was quantified, sonicated, denatured and transferred onto Nylon membrane (RPN303B, GE Healthcare) according to the Cell Signaling DNA Dot Blot Protocol. 5-methylcytosine (5-mC) Ab (#28692, Cell Signaling) was used to detect global DNA methylation of the samples. Methylene blue staining was used to verify equal DNA loading across samples.

**Chromatin profiling**

Approximately 50,000 cells were harvested for ATAC-seq for each replicate. ATAC-seq was performed as previously described [23]. ATAC-seq libraries were sequenced with the Illumina HiSeq4000 using 100bp paired-end single indexed run. Raw reads were first QCed (*FASTQC*) and quality and adapter trimmed using *Trimmomatic*. Trimmed reads were then aligned to hg19 build of the human genome using Bowtie2 (v2.2.5) with alignment parameters: bowtie2 -X 2000– local–dovetail. Potential PCR duplicate reads were removed using *MarkDuplicates* from the *Picard* tools. Peaks were called in each sample using *MACS2* and further filtered using the ENCODE consensus blacklist regions (http://hgdownload.cse.ucsc.edu/goldenPath/hg19/encodeDCC/wgEncodeMapability/). Differential peaks were identified using R package *diffbind* across the consolidated peak sets and metrics such as adjusted p values were reported.

Wu et al
Supplemental Figure 1

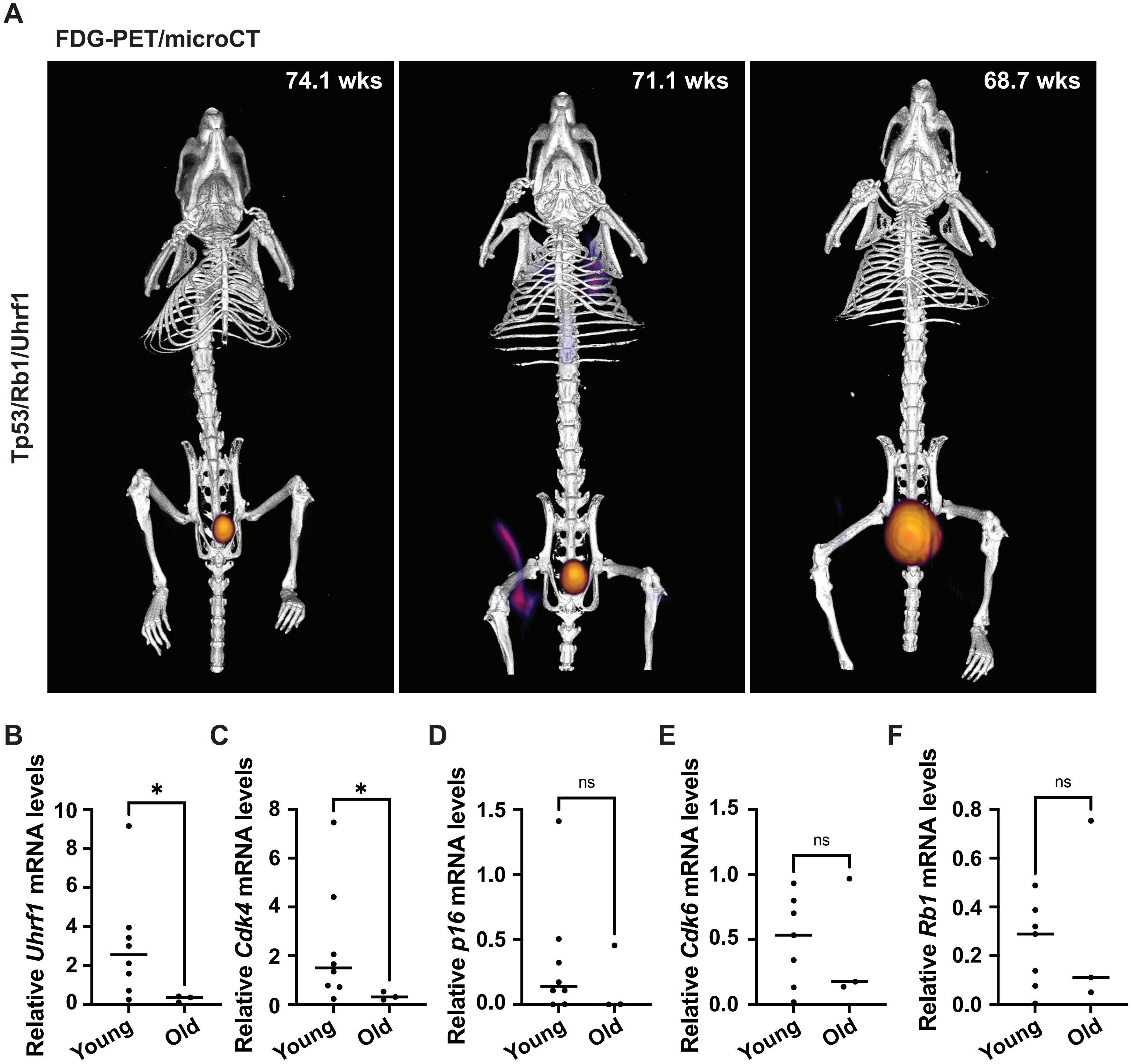

**Wu et al**
**Supplemental Figure 2**

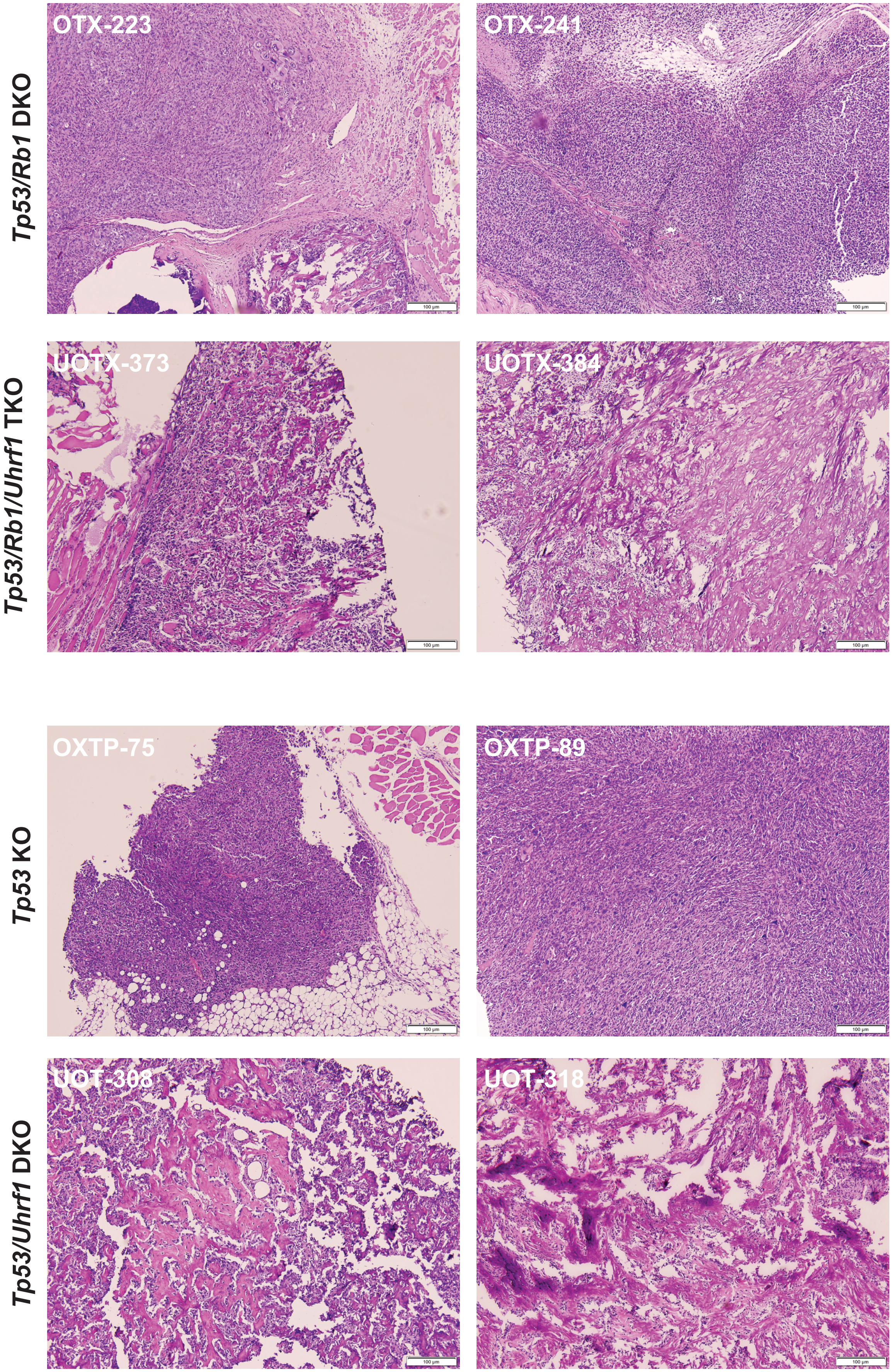

**Wu et al**
**Supplemental Figure 3**

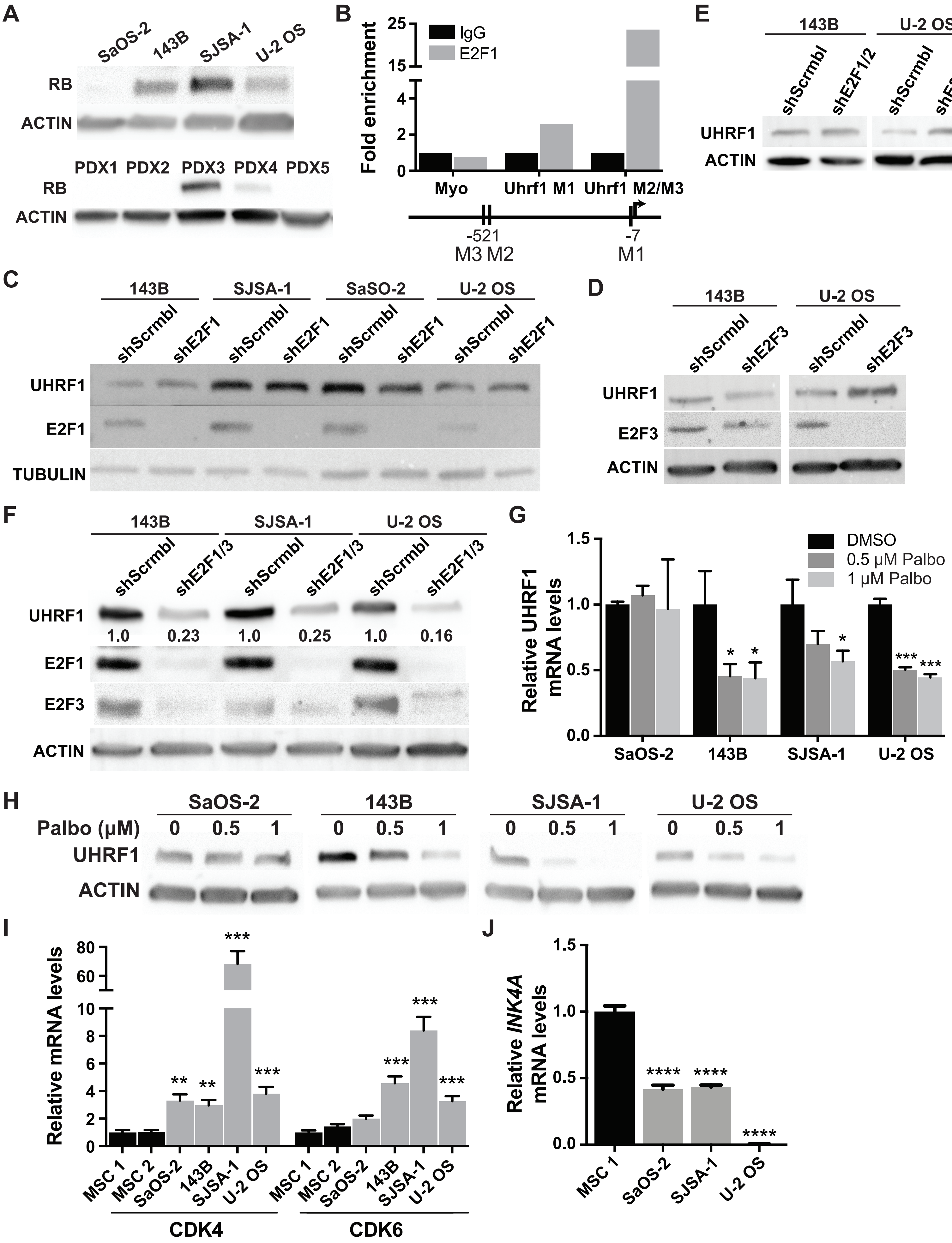

Wu et al
Supplemental Figure 4

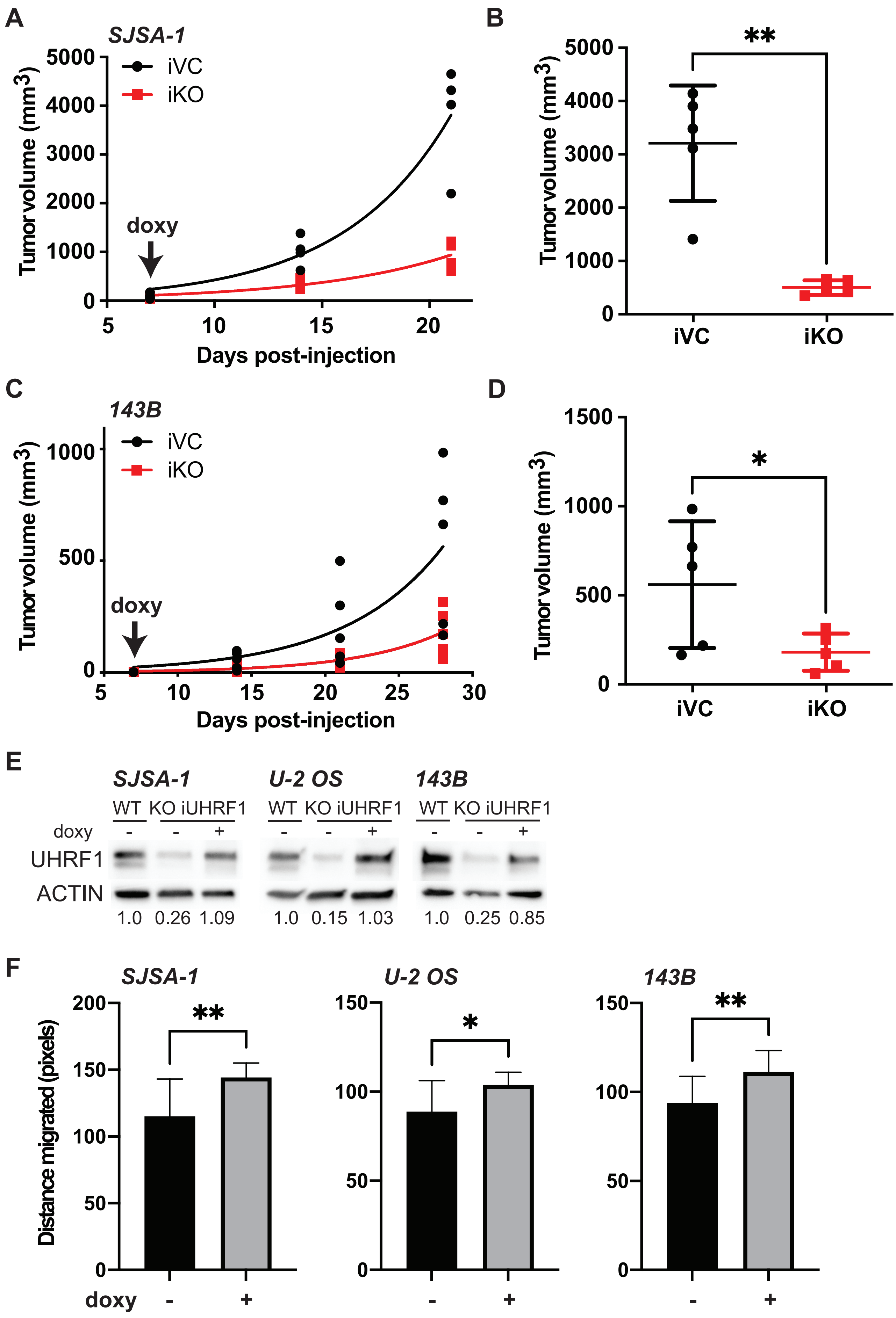

**Wu et al**
**Supplemental Figure 5**

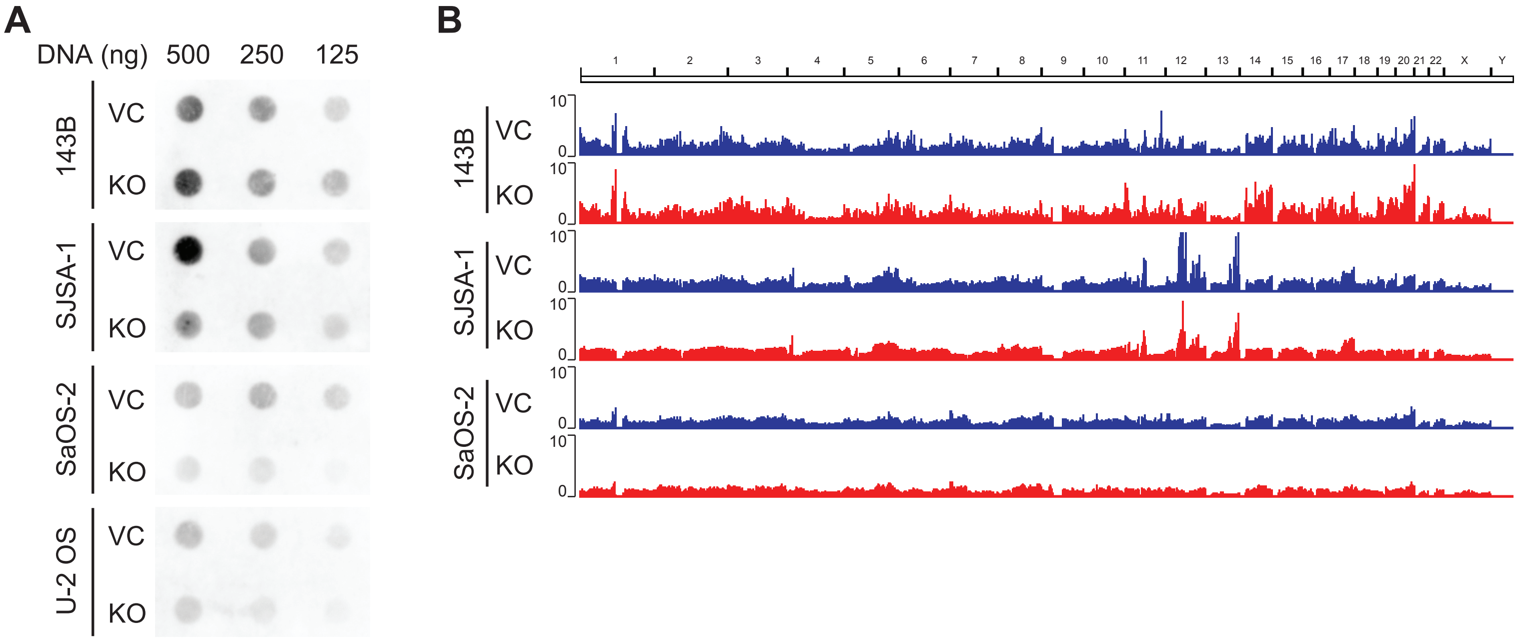

**Supplemental Table 1.** Pathology analysis of mouse osteosarcoma strains

| Genotype | Tumor ID | Malignant osteoid formation | Tumor nuclear size | Anaplasia/Significant pleomorphism | Mitotic activity | Necrosis |
|---|---|---|---|---|---|---|
| **p53 cKO** | OXTP75 | No | Medium to large sized with epithelioid features | Focal | 3/10 HPF | Yes, small focus |
| | OXTP89 | No | Medium to large sized | Yes | 3/10 HPF | No |
| | OXTP73 | Yes, abundant | Medium to large sized | No | 1/10 HPF | No |
| **p53/Rb1 DKO** | OTP223 | Yes, rare | Medium to large sized with epithelioid features | Yes | 4/10 HPF | No |
| | OTP196 | Yes, abundant | Medium sized, frequent MNT cells | No | 1-2/10 HPF | No |
| | OTP241 | No | Small to medium sized | Yes | 10/10 HPF | Yes |
| **p53/Uhrf1 DKO** | UOTX414 | Yes, abundant | Small sized | No | 0/10 HPF | No |
| | UOTX373 | Yes, abundant | small to Medium sized, rare MNT cells | No | 0/10 HPF | No |
| | UOTX384 | Yes, abundant | small to Medium sized, rare MNT cells | No | 1/10 HPF | No |
| **p53/Rb1/Uhrf1 TKO** | UOT259 | Yes, abundant | Medium sized, frequent MNT cells | No | 1/10 HPF | No |
| | UOT318 | Yes, abundant | Medium sized, occasional MNT cells | No | 1/10 HPF | No |
| | UOT308 | Yes, abundant | Medium sized, occasional MNT cells | No | 1/10 HPF | No |

**Supplemental Table 2.** ATAC-seq results p<0.05

| | seqnames | start | end | Conc_VO | Conc_UHRF | Fold | p.value |
|---|---|---|---|---|---|---|---|
| 7763 | chr5 | 38147946 | 38148446 | 3.36 | 6.07 | -2.71 | 0.000352 |
| 3317 | chr15 | 33406160 | 33406660 | 4.57 | 6.72 | -2.14 | 0.0129 |
| 7767 | chr5 | 38756061 | 38756561 | 4.92 | 6.76 | -1.85 | 0.0249 |
| 4377 | chr17 | 68223297 | 68223797 | 4.56 | 6.3 | -1.74 | 0.015 |
| 8608 | chr6 | 44084322 | 44084822 | 4.31 | 5.99 | -1.67 | 0.0285 |
| 5989 | chr20 | 23125981 | 23126481 | 4.16 | 5.77 | -1.62 | 0.0167 |
| 5965 | chr20 | 17519102 | 17519602 | 3.91 | 5.45 | -1.54 | 0.0138 |
| 5084 | chr2 | 23637633 | 23638133 | 5.18 | 6.5 | -1.31 | 0.0293 |
| 5213 | chr2 | 49056145 | 49056645 | 4.48 | 5.78 | -1.3 | 0.0492 |
| 4829 | chr19 | 30369822 | 30370322 | 5.99 | 7.24 | -1.25 | 0.034 |
| 6767 | chr3 | 46688121 | 46688621 | 4.46 | 5.66 | -1.2 | 0.0402 |
| 2655 | chr13 | 36705232 | 36705732 | 6.35 | 5.38 | 0.97 | 0.0439 |
| 9982 | chr9 | 4741018 | 4741518 | 7.73 | 6.66 | 1.07 | 0.0374 |
| 1176 | chr10 | 63422386 | 63422886 | 6.64 | 5.38 | 1.26 | 0.0303 |
| 9071 | chr7 | 25990604 | 25991104 | 6.38 | 4.93 | 1.45 | 0.0352 |
| 5460 | chr2 | 125593939 | 125594439 | 6.54 | 4.93 | 1.61 | 0.0328 |
